# Restructuring of amygdala subregion apportion across adolescence

**DOI:** 10.1101/690875

**Authors:** Claire E. Campbell, Adam F. Mezher, Sandrah P. Eckel, J. Michael Tyszka, Wolfgang M. Pauli, Bonnie J. Nagel, Megan M. Herting

## Abstract

Total amygdala volumes develop in association with sex and puberty, and postmortem studies find neuronal numbers increase in a nuclei specific fashion across development. Thus, amygdala subregions and composition may evolve with age. Our goal was to examine if amygdala subregion absolute volumes and/or relative proportion varies as a function of age, sex, or puberty in a large sample of typically developing adolescents (N=408, 43% female, 10-17 years). Utilizing the *in vivo* CIT168 atlas, we quantified 9 subregions and implemented Generalized Additive Mixed Models to capture potential non-linear associations with age and pubertal status between sexes. Only males showed significant age associations with the basolateral ventral and paralaminar subdivision (BLVPL), central nucleus (CEN), and amygdala transition area (ATA). Again, only males showed relative differences in the *proportion* of the BLVPL, CEN, ATA, along with lateral (LA) and amygdalostriatal transition area (ASTA), with age. Using a best-fit modeling approach, age, and not puberty, was found to drive these associations. The results suggest that amygdala subregions show unique developmental patterns with age in males across adolescence. Future research is warranted to determine if our findings may contribute to sex differences in mental health that emerge across adolescence.

## Introduction

The amygdala is a collection of nuclei located in the medial temporal lobe, with extensive connections to the cerebral cortex (Amaral & Price, 1984; Barbas & De Olmos, 1990; Ghashghaei & Barbas, 2002). The heterogeneous structure and function of the amygdala nuclei play a vital role in mediating a number of cognitive, affective, and motivational processes (Baxter & Murray, 2002; Bzdok, Laird, Zilles, Fox, & Eickhoff, 2013; Hariri, Tessitore, Mattay, Fera, & Weinberger, 2002; Meyer-Lindenberg et al., 2005; Raznahan et al., 2011; Tottenham & Gabard-Durnam, 2017). Cytoarchitecture and lesion studies have helped determine how these diverse groupings of amygdala neurons mediate specific processes (Amaral & Price, 1984; Amunts et al., 2005; Ghashghaei & Barbas, 2002; Krettek & Price, 1978; Solano-Castiella et al., 2011). Previous studies have shown the basal and lateral nuclei process high-level sensory input and emotional regulation (Sananes & Davis, 1992; Schoenbaum, Chiba, & Gallagher, 1999; Wan & Swerdlow, 1997), while the central nucleus is involved in reward and aversive learning (Baxter & Murray, 2002; Killcross, Robbins, & Everitt, 1997). Moreover, the paralaminar nucleus of the amygdala contains neurons that continue to mature and migrate into adulthood (Amaral & Price, 1984; Bernier, Bedard, Vinet, Levesque, & Parent, 2002; deCampo & Fudge, 2012; Tosevski et al., 2002); this region’s potential for regional neural plasticity (deCampo & Fudge, 2012) may be involved in modulating changes in nuclei size and thus the restructuring of amygdala composition.

When treating the amygdala as a singular unit, total amygdala volumes continue to increase from childhood to young adulthood, with distinct developmental patterns seen based on sex and pubertal stage (Bramen et al., 2011; Giedd et al., 1996; M. M. Herting et al., 2014; M. M. Herting et al., 2018; L. Wierenga et al., 2014; L. M. Wierenga et al., 2018). However, a recent postmortem study (N=24 neurotypical brains, ages 2-48 years) found that neuron numbers increase in the amygdala but do so in a nucleus specific manner (Avino et al., 2018). These findings suggest that regionally specific neuronal increases may contribute to age related differences in amygdala subregion volumes and/or subregional apportionment across childhood and adolescence. Accordingly, our study aimed to quantify both the absolute volumes as well as the relative proportion, or the *relative volume fraction (RVF)*, of amygdala subregions in a large cross-sectional sample of 408 children and adolescents (n=177 females, ages 10-17 years) as well as examine if these patterns varied as a function of age, sex, or pubertal status. While previous amygdala atlases have been limited as they have been created by *ex vivo* brains from small samples of elderly, male individuals (Amunts et al., 2005; Saygin et al., 2017), we implemented a newly developed high-resolution probabilistic atlas, known as the CIT168, which was created using *in vivo* MRI data from 168 (50% female) healthy young adult brains (Pauli, Nili, & Tyszka, 2018; Tyszka & Pauli, 2016). Using this approach, we segmented the amygdala into nine bilateral regions of interest (ROIs), including the lateral nucleus (LA), basolateral dorsal and intermediate subdivision (BLDI), basolateral ventral and paralaminar subdivision (BLVPL), basomedial nucleus (BM), cortical and medial nuclei (CMN), central nucleus (CEN), anterior amygdala area (AAA), amygdala transition areas (ATA), and amygdalostriatal transition area (ASTA) (**Table 1**). Building on previous research (Baxter & Murray, 2002; M. M. Herting et al., 2014; M. M. Herting et al., 2018; Janak & Tye, 2015; Rollins & King, 2000; Tyszka & Pauli, 2016; L. Wierenga et al., 2014; L. M. Wierenga et al., 2018), we then examined how age, sex, and pubertal status were associated with subregion volumes as well as the relative proportion of each subregion within the amygdala in adolescents. Given that the basolateral nucleus increases innervation with the prefrontal cortex during adolescent neurodevelopment (Cunningham, Bhattacharyya, & Benes, 2002) and the paralaminar’s potential for postnatal neuroplasticity (deCampo & Fudge, 2012), we hypothesized that lateral, basal, and paralaminar subregions would be larger as a function of age across adolescence, as well as occupy a larger proportion of the amygdala with age. Ultimately, understanding how the human amygdala develops throughout adolescence may help discern developmental changes seen in social-emotional and reward-related behavior, as well as identify risk factors for mental health disorders.

**Table 1.** Sample characteristics.

*A) Demographics of study participants*
|  | All |  |  | Female |  |  | Male |  |  | Difference between Male and Female |  |  |
| --- | --- | --- | --- | --- | --- | --- | --- | --- | --- | --- | --- | --- |
|  | N | Mean | SD | N | Mean | SD | N | Mean | SD | df | Coefficient | p-value |
| Age | 408 | 14.12 | 1.62 | 177 | 14.03 | 1.57 | 231 | 14.19 | 1.66 | 404 | 0.14 | 0.93 |
| BMIz | 408 | 0.47 | 0.96 | 177 | 0.45 | 0.90 | 231 | 0.48 | 1.01 | 404 | 0.37 | 0.55 |
| ICV | 408 | 1468317 | 137346 | 177 | 1377606 | 101538 | 231 | 1537823 | 119612 | 404 | 117942 | **<0.0001** |
| PDS | 408 | 3 | 1 | 177 | 3 | 1 | 231 | 3 | 1 | 404 | -0.22 | 0.745 |
| SES | 408 | 27.94 | 13.67 | 177 | 28.46 | 14.53 | 231 | 27.53 | 12.99 | 404 | 0.48 | 0.09 |

*B) Associations between predictors*
|  | All |  |  | Female |  |  | Male |  |  | Difference between Male and Female |  |  |
| --- | --- | --- | --- | --- | --- | --- | --- | --- | --- | --- | --- | --- |
|  | df | Coefficient | p-value | df | Coefficient | p-value | df | Coefficient | p-value | df | Coefficient | p-value |
| Age to ICV | 406 | 4.5E-07 | 0.45 | 175 | 1.7E-07 | 0.88 | 229 | 1.3E-07 | 0.89 | 404 | -3.0E-08 | 0.98 |
| ICV to BMIz | 406 | -7679 | 0.28 | 175 | -534 | 0.95 | 229 | -14206 | 0.07 | 404 | -9668 | 0.25 |
| BMIz to Age | 406 | 0.00 | 0.95 | 175 | 0.02 | 0.68 | 229 | -0.016 | 0.69 | 404 | -0.02 | 0.57 |
| PDS to ICV | 406 | -1.1E-06 | **0.0001** | 175 | 1.5E-07 | 0.77 | 229 | -1.6E-09 | 1.00 | 404 | -1.1E-07 | 0.82 |
| BMIz to PDS | 406 | 0.12 | 0.05 | 175 | 2.6E-01 | **0.006** | 229 | 6.3E-02 | 0.49 | 404 | -1.4E-01 | 0.14 |
| SES to Age | 406 | -0.14 | 0.73 | 175 | -0.01 | 0.98 | 229 | -0.21 | 0.69 | 404 | -0.14 | 0.82 |
| SES to PDS | 406 | 1.41 | 0.11 | 175 | 1.70 | 0.28 | 229 | 1.13 | 0.33 | 404 | -4.1E-01 | 0.76 |
| SES to BMIz | 406 | 1.35 | 0.05 | 175 | 1.84 | 0.13 | 229 | 1.06 | 0.21 | 404 | -0.55 | 0.59 |

Notes: P-values with significance level of less than 0.05 are bolded. Abbreviations: SD, standard deviation, BMIz, Body Mass Index z-score; PDS, Pubertal Development Scale; ICV, Intracranial Volume.

## Materials and Methods

### Participants and Measures

This study incorporated cross-sectional data from 408 adolescents (n=177 females), ages 10 to 17 years (**Supporting Figure 1**), from ongoing research studies at Oregon Health & Science University. A comprehensive telephone interview was conducted to determine eligibility for all participants, and written consent and assent were obtained from each participating adolescent and at least one of their biological parents. All participants were right-handed and free of neurological, neurodevelopmental, and/or psychological diagnoses based on the DISC Predictive Scales (Shaffer, Fisher, Lucas, Dulcan, & Schwab-Stone, 2000). Subjects were also excluded if they: obtained head trauma; had serious medical problems; reported learning disabilities; used any medications that affected their central nervous system; consumed >10 lifetime alcoholic drinks, >2 alcoholic drinks on one occasion, >10 lifetime uses of marijuana, >4 lifetime uses of cigarettes, and any other drug use history; had maternal alcohol use of 4 drinks per occasion or 7 drinks per week or other drugs during pregnancy; parental history of bipolar I or psychotic disorders; inadequate English skills; sensory issues; non-removable mental in their body. The prior exclusionary criteria can be found in greater detail in cited papers (Alarcon, Cservenka, Rudolph, Fair, & Nagel, 2015; Morales, Jones, Ehlers, Lavine, & Nagel, 2018; Scheuer et al., 2017).

Based on prior research (Baxter & Murray, 2002; M. M. Herting et al., 2014; M. M. Herting et al., 2018; Janak & Tye, 2015; Rollins & King, 2000; Tyszka & Pauli, 2016; L. Wierenga et al., 2014; L. M. Wierenga et al., 2018), we considered three primary biological and physical factors for each participant: age, sex, and pubertal status. Pubertal status was determined by self-report using the Pubertal Development Scale (PDS) (Petersen, Crockett, Richards, & Boxer, 1988), with scores for each of the 5 questions ranging from 1 (not started) to 4 (development seems complete). Scores across the items were averaged to a single comprehensive score. Based on the literature, we also identified two important covariates, including social economic status (SES) (Brito & Noble, 2014) and body mass index (BMI) (Perlaki et al., 2018). SES was assessed utilizing the Hollingshead Four-Factor Index of Socioeconomic Status-Child (Hollingshead, 1975) and BMI using the Centers for Disease Control and Prevention’s BMI Percentile Calculator for Child and Teen English Version (http://nccd.cdc.gov/dnpabmi/Calculator.aspx) by providing participant birth date, date of measurement, sex, height (to nearest 0.1 cm) and weight (to nearest 0.1 kg); subjects were removed if they did not have BMI values (**Supporting Figure 2**). BMI z-scores (BMIz), which correspond to growth chart percentiles, were then calculated to reflect the relative weight of the individual using the appropriate reference standard based on the individual’s age and sex (Must & Anderson, 2006).

### MRI Data Collection and Preprocessing

A whole-brain T1-weighted MRI scan was acquired for each participant on the same 3 Tesla MRI system (Magnetom Tim Trio, Siemens Medical Solutions, Erlangen, Germany) using a 12-channel head coil at the Oregon Health & Science University’s Advanced Imaging Research Center (TR = 2300ms, TE = 3.58ms, TI = 900ms, flip angle = 10°, 256×240 matrix, voxel size = 1.0 mm × 1.0 mm × 1.1 mm). Raw images were quality checked for motion and given a rating of 1 (pass), 2 (review), or 3 (fail) (Backhausen et al., 2016); for details, see **Supporting Figure 2**. Using the Functional Magnetic Resonance Imaging of the Brain Software Library (FSL) version 5.0 (Jenkinson, Beckmann, Behrens, Woolrich, & Smith, 2012; Smith et al., 2004; Woolrich et al., 2009). Each brain image was first reoriented to standard orientation using FSL’s *fslreorient2std* function. Images were then automatically cropped to reduce lower head and neck using FSL’s *robustfov* tool and rigid-body AC-PC aligned. Using the *antsBrainExtraction* function from the Advanced Normalization Tools (ANTs, Version 2.1.0.post691-g9bc18) (B. B. Avants et al., 2011), each image was skull-stripped to allow for an N4 Bias Field Correction (Tustison et al., 2010) on the whole-brain image.

### Probabilistic Amygdala Volumes and Relative Volume Fractions (RVF)

Details of the *in vivo* amygdala probabilistic atlas construction, validation, estimates of individual differences, and comparison with previous atlas’ have been previously reported (Pauli et al., 2018; Tyszka & Pauli, 2016). As previously published (Megan M Herting et al., 2020), we adapted this technique to allow for each youth’s image to be registered to the CIT168 atlas using a B-spline bivariate symmetric normalization (SyN) diffeomorphic registration algorithm from ANTs version 2.2 (B. Avants, Anderson, Grossman, & Gee, 2007). Applying the inverse diffeomorphism mapped the CIT168 probabilistic atlas labels to the individual space of each participant, yielding probabilistic ROIs for left and right total amygdala and the following nine subregions: lateral nucleus (LA); dorsal and intermediate divisions of the basolateral nucleus (BLDI); ventral division of the basolateral nucleus and paralaminar nucleus (BLVPL); basomedial nucleus (BM); central nucleus (CEN); cortical and medial nuclei (CMN); amygdala transition areas (ATA); amygdalostriatal transition area (ASTA); and anterior amygdala area (AAA). Descriptions of each subregion can be found in **Supporting Table 1**. The quality of all amygdala segmentations was confirmed visually (A.F.M.). Total ROI volumes were estimated by summation over the entire brain of voxel volumes weighted by the probability that a voxel belonged to the ROI. This approach differs from deterministic voxel summation following application of an arbitrary probability threshold (typically p > 0.5) in that uncertainty about the true extent of the ROI continues to be accurately represented in the resulting volume estimate. In addition to the absolute volume of the whole amygdala and each ROI generated from the CIT168 atlas, the fractional volume of each ROI within the amygdala was estimated relative to the total amygdala volume in each hemisphere, referred to subsequently as the relative volume fraction (RVF) for a given ROI.

Although the CIT168 atlas labels were delineated in high CNR template images, the amygdala nuclei in individual T1-weighted images have much lower CNR. The accuracy of volume estimates of amygdala subregions is heavily dependent on the accuracy of the SyN diffeomorphic registration between the high-CNR CIT168 space and the low-CNR individual space, where subnuclear boundaries become very indistinct to a human observer when viewed in 2D sections. Prior work modeling the accuracy of volume estimates in low CNR scenarios (Tyszka and Pauli (2016) suggests the implementation of the SyN diffeomorphic algorithm used in this study is relatively robust at CNRs of 1.0 and higher. This robustness is likely a consequence of the registration cost function drawing information from a local volume (approximately 4 voxel radius) to optimize the local deformation. Consequently, any subject with an intra-amygdala CNR less than 1.0 in either the right or left amygdala was excluded from further analyses (**Supporting Figure 2**). Intra-amygdala CNR was estimated as follows: 1. The intensity contrast within the amygdala was estimated from the interquartile range (IQR) of intensities within the entire amygdala from each subject’s T1-weighted image. 2. The standard deviation (SD) of the noise was estimated from the residual signal obtained when the T1-weighted atlas template image was subtracted from each subject’s T1-weighted image. 3. The CNR was estimated as the ratio of the IQR to noise SD within the amygdala. The group mean SD over all subjects of residual signal within the amygdala was 24 for the right and 25 for the left hemisphere. The mean lower and upper quartile intensities within the amygdala were 278 and 311 (IQR = 33) for the right hemisphere and 277 and 310 (IQR = 33) for the left hemisphere. The average CNR was 1.4 for the amygdala in both hemispheres in our sample, suggesting the current study has sufficient CNR necessary to support robust amygdala subregion volume estimation when using SyN diffeomorphic registration (Tyszka & Pauli, 2016).

A 2-Dimensional visual representation of the probabilistic maps of each amygdala subregion segmentation on a representative subject can be seen in **Figure 1**. To fully demonstrate the CIT168 segmentation, overlay images of coronal slices through the entire rostral-caudal extent of the amygdala with boundary outlines for each ROI is also presented for four subjects in **Figure 2**. The subjects were randomly chosen to cover the distributions of our age range, including 1 male and 1 female from both the early and older adolescent periods. We also provide probabilistic estimates at the group level for a subset (N=52) of our sample in **Supporting Figure 3**.

**Figure 1:**
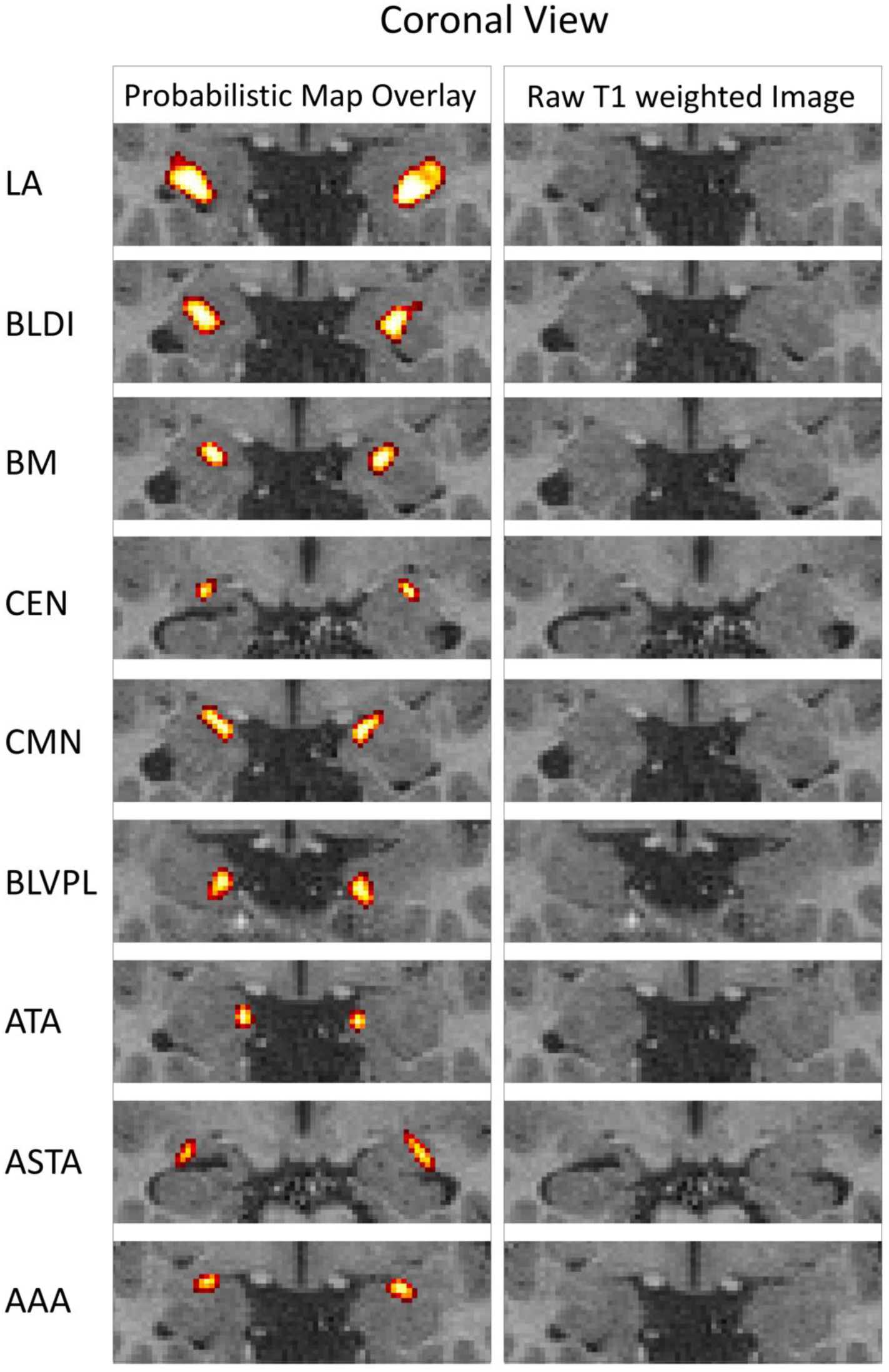
Probabilistic map of each amygdala subregion in a representative adolescent. Structural MRI with the left column showing the probabilistic maps of the 9 bilateral subregions shown in the coronal view (thresholded at probabilistic value of .15 for visualization purposes) and showing the raw T1 weighted image in the same coronal slice. Key: LA, lateral nucleus; BLDI, basolateral dorsal and intermediate subdivision; BLVPL, basolateral ventral and paralaminar subdivision; BM, basomedial nucleus; CMN, cortical and medial nuclei; CEN, central nucleus; AAA, anterior amygdala area; ATA, amygdala transition area; ASTA, amygdalostriatal transition area.

**Figure 2:**
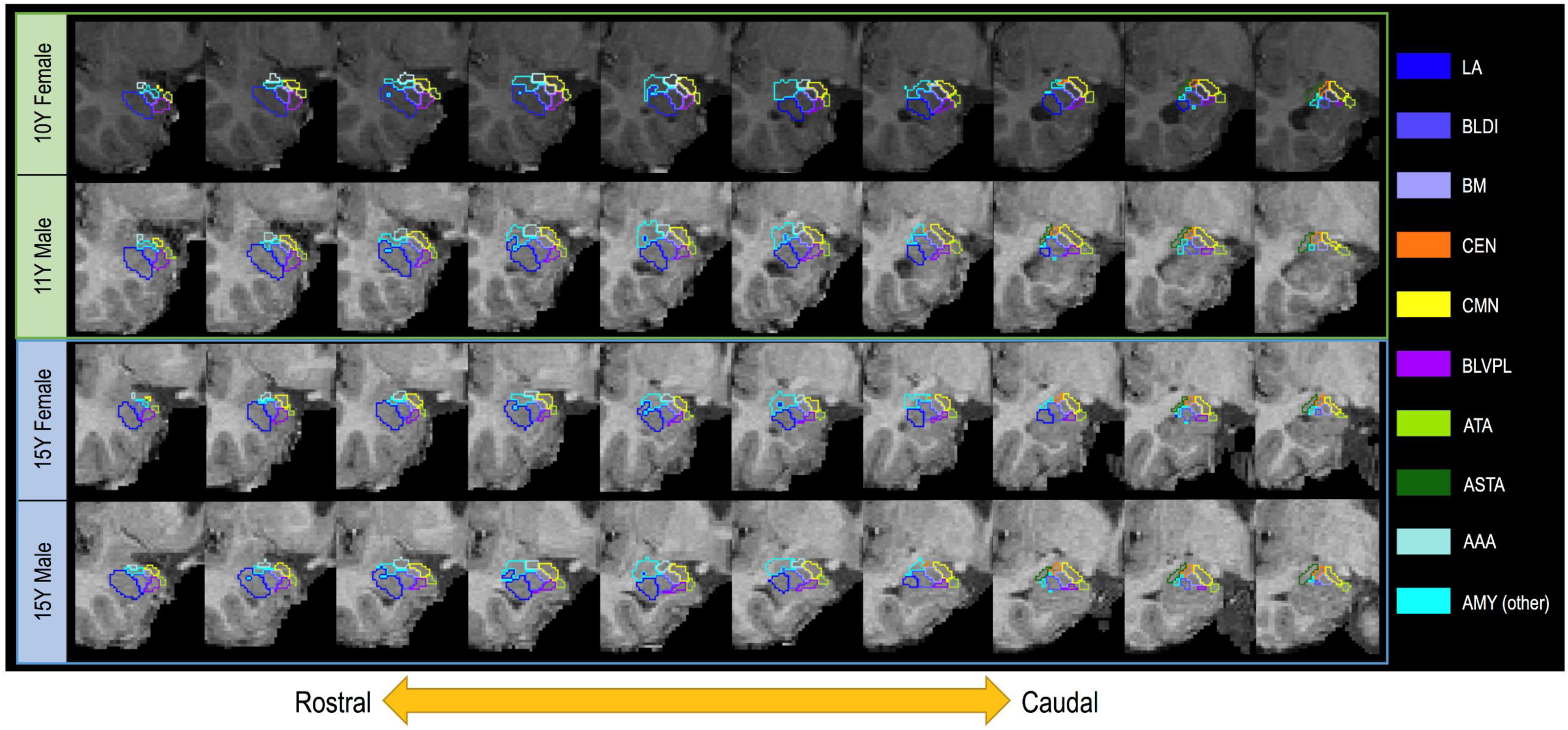
Outline of CIT168 segmentation on coronal slices through entire rostral-caudal view of the amygdala in the right hemisphere for four representative subjects. A maximum likelihood label was created for each subregion of the amygdala by creating a label based on a simple competition between probabilistic labels with a thresholded probabilistic value of .3 for visualization purposes; slices (1mm) are sequential (no gap).

### Statistical Analysis

Data were analyzed in R (version 3.5.1). Linear regressions (M1) were utilized to examine the associations between independent variables and covariates, including age and intracranial volume (ICV), ICV and BMIz, BMIz and age, PDS to ICV, BMIz to PDS, and SES to age. These associations were also assessed across all participants and between males and females to see if the associations were significantly different by sex: 

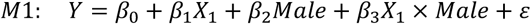

To examine if absolute probabilistic volumes and composition (i.e. proportional volumes or RVFs) related to age, sex, and pubertal status, we employed a Generalized Additive Mixed Model (GAMM) implemented by the *mgcv* package (version 1.8-24 in R version 3.5.1, R Core Team, 2018). Given that this developmental period shows non-linear subcortical brain volume growth patterns (M. M. Herting et al., 2018; L. Wierenga et al., 2014), a GAMM approach was chosen as it allows for data-driven estimation of non-linear associations (with linearity as a special case), using ‘smooth’ functions, s(), in place of linear terms. To examine the association between age and amygdala probabilistic volumes and composition (RVF), as well as determine if these associations vary by sex, each ROI was modeled independently using a GAMM (M2/M3) with fixed effects including smooth terms for age and age-by-sex (s_1_ and s_2_, respectively), as well as a linear term for sex, hemisphere, BMIz, ICV (for absolute volumes), SES, and a random intercept (*U*_*i*_) for participant *i*: 

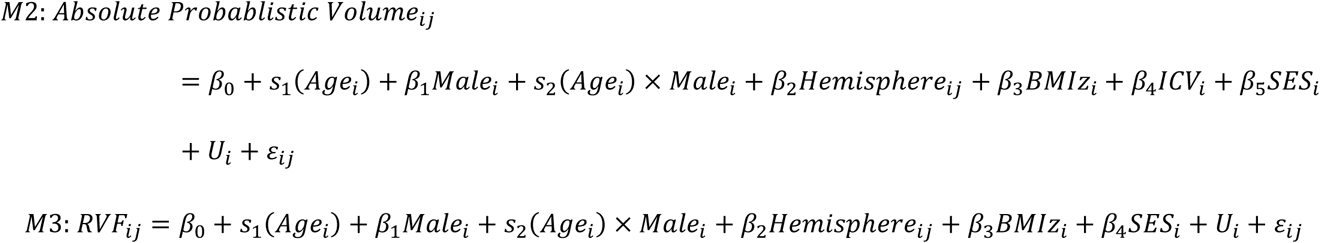

Where *Absolute Probablistic Volume*_*i,j*_ and RVF_*i,j*_ were defined for each subject, *i*, in either the left or right hemisphere, *j*. Hemisphere was a nested term – for left and right – accounting for the fact that hemispheres from the same subject will be more similar than hemispheres between subjects. Each smooth term is a shrinkage version of a cubic regression spline with four equally spaced knots. ICV, SES, and BMIz are added as covariates. SES and BMI has been shown to relate to amygdala volumetric differences during adolescence (Brito & Noble, 2014; Perlaki et al., 2018). ICV accounts for the varying volumes of brain regions given the head size of the individual and is added to the absolute probabilistic volume models. For RVF, ICV was not added since the effect of ICV would likely not provide a significant influence given the inherent control when creating a fraction with the total amygdala.

Given that markers of pubertal development have been shown to relate to total amygdala volumes across adolescence (Goddings et al., 2014; M. M. Herting et al., 2014; L. M. Wierenga et al., 2018), we then utilized a model building strategy to determine if age, pubertal development, or their combination best predicted amygdala subregion absolute probabilistic volumes and RVFs across adolescence. Given that pubertal development follows a different age-related trajectory in males versus females and physical changes are distinct in males (e.g. facial hair, testes development) and females (e.g. breast development, menstruation) (Berenbaum, Beltz, & Corley, 2015), these analyses were performed in each sex separately. First, in each sex we examined the smooth effect of age (M4/M7). Next, we examined the smooth effect of pubertal stage (M5/M8). Lastly, we examined both the smooth effects of age and pubertal stage as well as the interaction term of age-by-pubertal stage (M6/M9), with smooths implemented by tensor product interactions, allowing for main effects and the interaction. Each model also included the fixed effects of BMIz, hemisphere, ICV (for absolute probabilistic volume models), and a random intercept (*U*_*i*_) for participant *i*: 

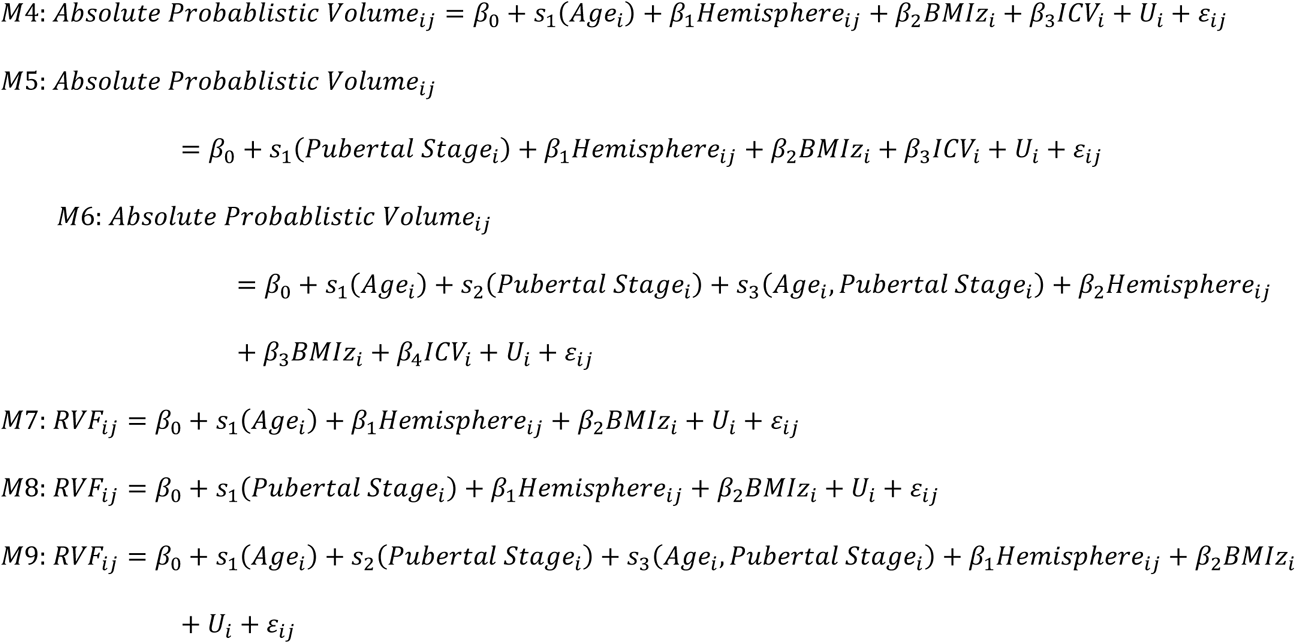

Akaike Information Criterion (AIC) and Likelihood ratio tests (p < 0.05) were used to compare model fits. To reduce type I error, each set of models across the 9 ROIs were corrected for multiple comparisons using the Bonferroni correction method (Bonferroni, 1936), with p-values <0.0056 deemed significant.

## Results

Males and females did not differ in age, BMIz, pubertal status (PDS), or socioeconomic status (SES), though on average, males had a significantly larger ICV compared to females (β=117942, p=<0.0001) (**Table 1A**). Associations between the variables did not significantly differ between the sexes (p’s>0.05) (**Table 1B**).

### Amygdala subregions: Age and sex effects

The mean and coefficient of variance (CoV) of the absolute probabilistic estimates and RVFs for each subregion using the CIT168 are summarized by hemisphere and sex in **Figure 3** and **Table 2**. From largest to smallest, absolute probabilistic estimates were on average 333-391 mm^3^ for the LA (∼21-22% of amygdala volume); 199-to 230 mm^3^ for the BLDI (∼12-13% of amygdala volume); 171-195 mm^3^ for the CMN (∼11% of the amygdala volume); 119-141 mm^3^ for the BLVPL (∼7-8% of the amygdala volume); 114 to 131 mm^3^ for the BM (∼7% of the amygdala volume); 93-111 mm^3^ for the ATA (∼6% of the amygdala volume); 69-77 mm^3^ for the ASTA (∼4 % of the amygdala volume); 63-71 mm^3^ for the AAA (∼4% of the amygdala volume); 47-53 mm^3^ for the CEN (∼3% of the amygdala volume); for total amygdala volume the range was 1582-1804 mm^3^.

**Table 2.** Probabilistic amygdala subregions by sex.

|  |  | LA |  | BLDI |  | BLVPL |  | BM |  | CMN |  | CEN |  | AAA |  | ATA |  | ASTA |  | Total |  |
| --- | --- | --- | --- | --- | --- | --- | --- | --- | --- | --- | --- | --- | --- | --- | --- | --- | --- | --- | --- | --- | --- |
| Sex | Hemisphere | Mean | CoV (%) | Mean | CoV | Mean | CoV | Mean | CoV | Mean | CoV | Mean | CoV | Mean | CoV | Mean | CoV | Mean | CoV | Mean | CoV |
| Female | Left | 333.78 | 9.73 | 202.35 | 9.71 | 122.42 | 10.69 | 115.67 | 9.74 | 172.01 | 8.96 | 48.19 | 9.94 | 63.08 | 10.23 | 93.11 | 9.93 | 69.83 | 10.47 | 1582.43 | 8.27 |
|  | Right | 343.82 | 11.00 | 199.02 | 10.54 | 119.55 | 10.87 | 114.52 | 10.05 | 171.35 | 9.22 | 47.17 | 10.36 | 62.75 | 11.24 | 96.78 | 10.12 | 69.31 | 11.22 | 1593.82 | 9.04 |
| Male | Left | 379.49 | 11.45 | 229.99 | 10.21 | 141.37 | 11.74 | 131.25 | 10.12 | 194.88 | 9.44 | 53.37 | 9.74 | 70.69 | 10.25 | 108.77 | 11.25 | 77.05 | 9.82 | 1796.00 | 8.72 |
|  | Right | 391.34 | 10.90 | 225.63 | 9.80 | 136.96 | 11.31 | 129.17 | 10.24 | 192.66 | 9.32 | 52.30 | 10.53 | 70.84 | 10.91 | 111.14 | 10.62 | 76.61 | 10.51 | 1804.39 | 8.60 |

*B) Relative volume fraction for each subregion (to total amygdala volume)*
|  |  | LA |  | BLDI |  | BLVPL |  | BM |  | CMN |  | CEN |  | AAA |  | ATA |  | ASTA |  |
| --- | --- | --- | --- | --- | --- | --- | --- | --- | --- | --- | --- | --- | --- | --- | --- | --- | --- | --- | --- |
| Sex | Hemisphere | Mean | CoV (%) | Mean | CoV | Mean | CoV | Mean | CoV | Mean | CoV | Mean | CoV | Mean | CoV | Mean | CoV | Mean | CoV |
| Female | Left | 0.21 | 4.92 | 0.13 | 2.78 | 0.08 | 5.45 | 0.07 | 3.80 | 0.11 | 4.65 | 0.03 | 6.88 | 0.04 | 6.07 | 0.06 | 6.69 | 0.04 | 6.80 |
|  | Right | 0.22 | 5.32 | 0.12 | 2.85 | 0.07 | 5.24 | 0.07 | 3.98 | 0.11 | 4.55 | 0.03 | 6.33 | 0.04 | 6.64 | 0.06 | 6.80 | 0.04 | 7.15 |
| Male | Left | 0.21 | 5.89 | 0.13 | 3.01 | 0.08 | 6.01 | 0.07 | 4.91 | 0.11 | 5.51 | 0.03 | 7.85 | 0.04 | 6.70 | 0.06 | 7.60 | 0.04 | 7.16 |
|  | Right | 0.22 | 5.79 | 0.12 | 2.87 | 0.08 | 6.08 | 0.07 | 4.79 | 0.11 | 4.97 | 0.03 | 7.78 | 0.04 | 6.80 | 0.06 | 7.65 | 0.04 | 7.46 |

**Figure 3:**
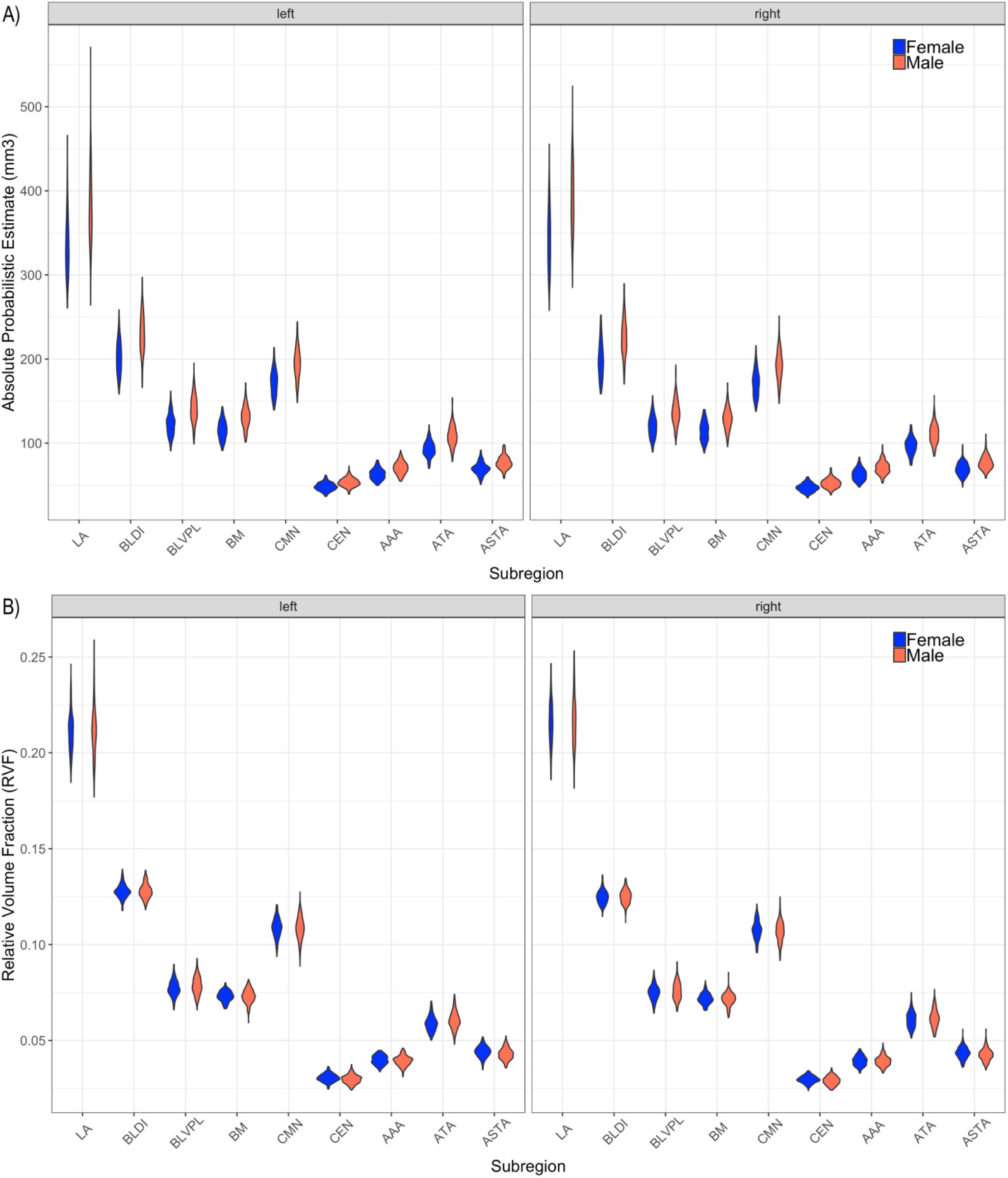
Amygdala subregion volumes and relative volume fractions in adolescent males and females. A) Probabilistic volumes (mm^3^) and B) relative volume fraction (RVF; proportional to total amygdala volume) for each of the 9 bilateral amygdala subregion ROIs.

GAMM model results examining the associations between total amygdala and each absolute probabilistic estimate (M2) or *proportion* (i.e. RVF) of the amygdala occupied by each subregion (M3) with age, sex, and age-by-sex interactions are presented in **Table 3** and **Table 4**, respectively. A significant age-by-sex interaction was detected for both the absolute probabilistic volumes for the BLVPL (Adj R^2^=.46), CEN (Adj R^2^=.45), and ATA (Adj R^2^=.52) (**Figure 4**), as well as the *proportion* of the amygdala occupied by these regions (BLVPL (Adj R^2^=.13), CEN (Adj R^2^=.09), ATA (Adj R^2^=.12)) (**Figure 5**). While trends were seen for the age-by-sex interaction for absolute volumes for the LA (p=0.095) and ASTA (p=0.04), age related differences in the *proportion* of the amygdala occupied by the LA (Adj R^2^=.06) and ASTA (Adj R^2^=.06) were significant between the sexes (**Figure 5**). Specifically, the LA, CEN, and ASTA were proportionately larger (as indexed by larger RVF values) with age, whereas the BLVPL and ATA were proportionately smaller with age in males. In contrast, no absolute or proportional differences were detected with age in females. A significant main effect of sex was also seen for the absolute volumes of the BLDI, BM, CMN, AAA, and Total Amygdala, with males having larger volumes even after accounting for differences in ICV (**Supporting Figure 4**). We also ensured findings of relative proportions were not driven by sex differences in ICV, as findings remained after including ICV in the RVF models (**Supporting Information** and **Supporting Table 2**).

**Table 3.**
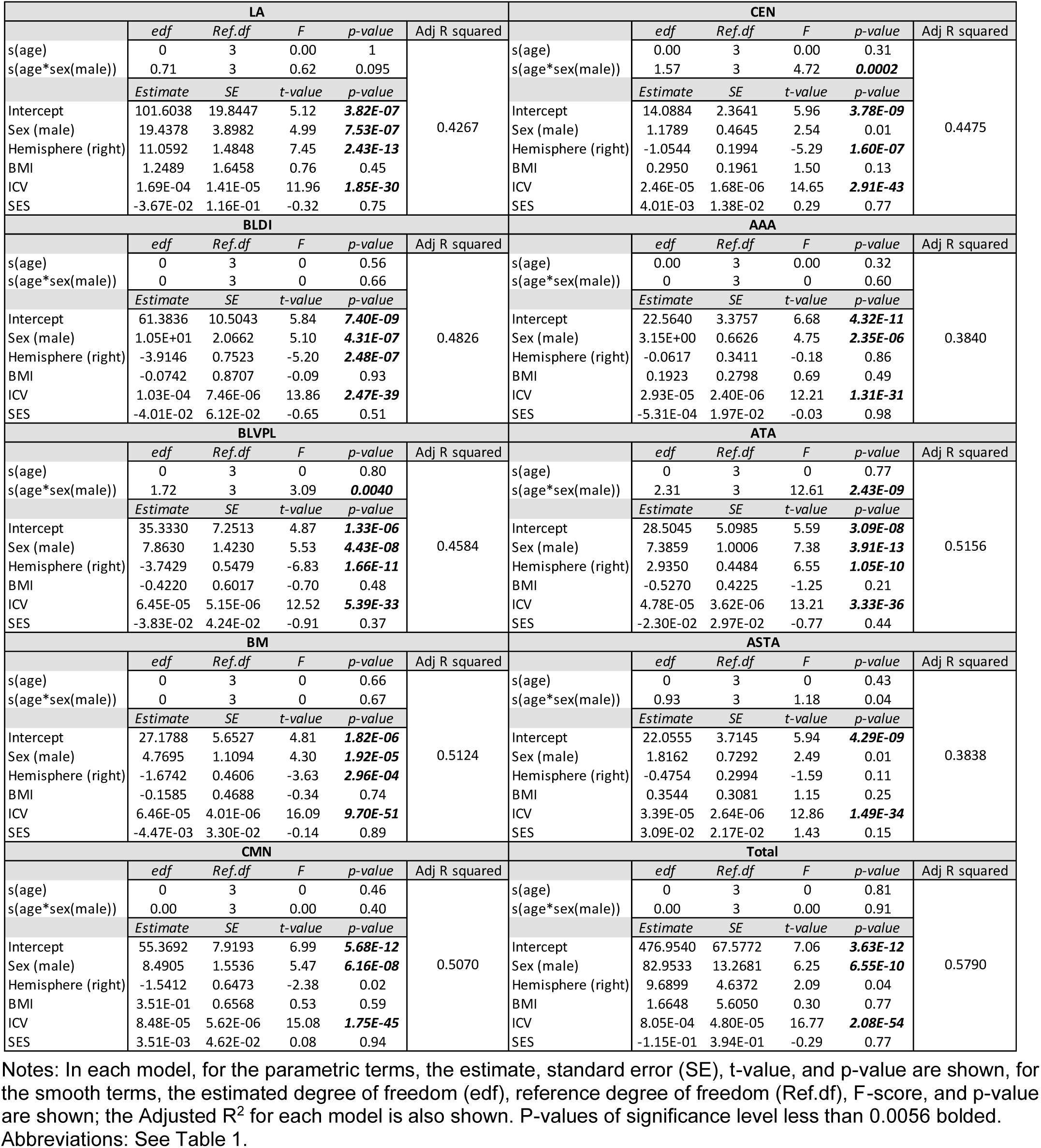
GAMM results for amygdala subregion Absolute Probabilistic Estimates associations with age, sex, and age sex interaction, controlling for hemisphere, BMI, and ICV.

**Table 4.**
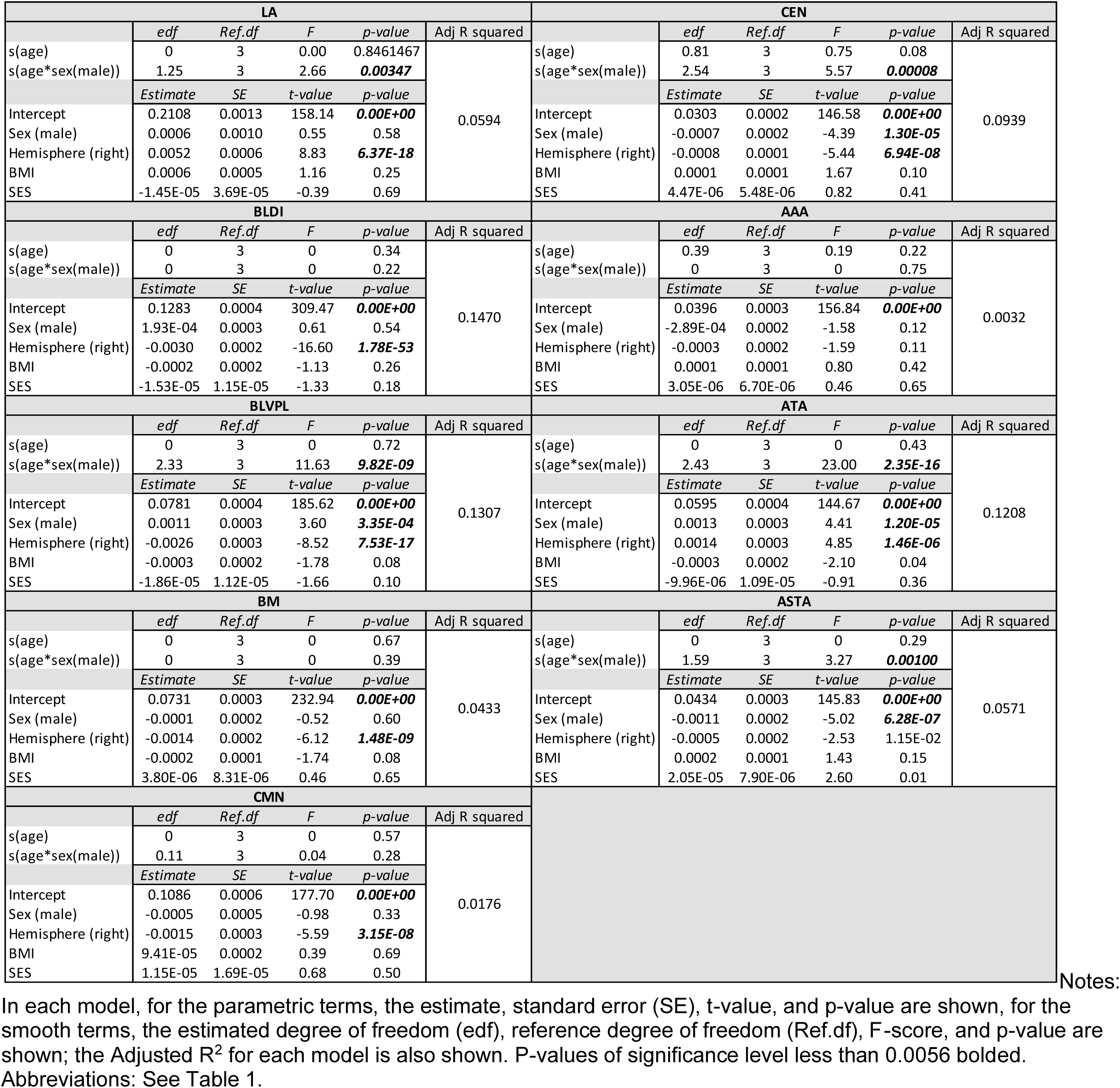
GAMM results for amygdala subregion Relative Volume Fraction (RVF) associations with age, sex, and age sex interaction, controlling for hemisphere and BMI.

**Figure 4:**
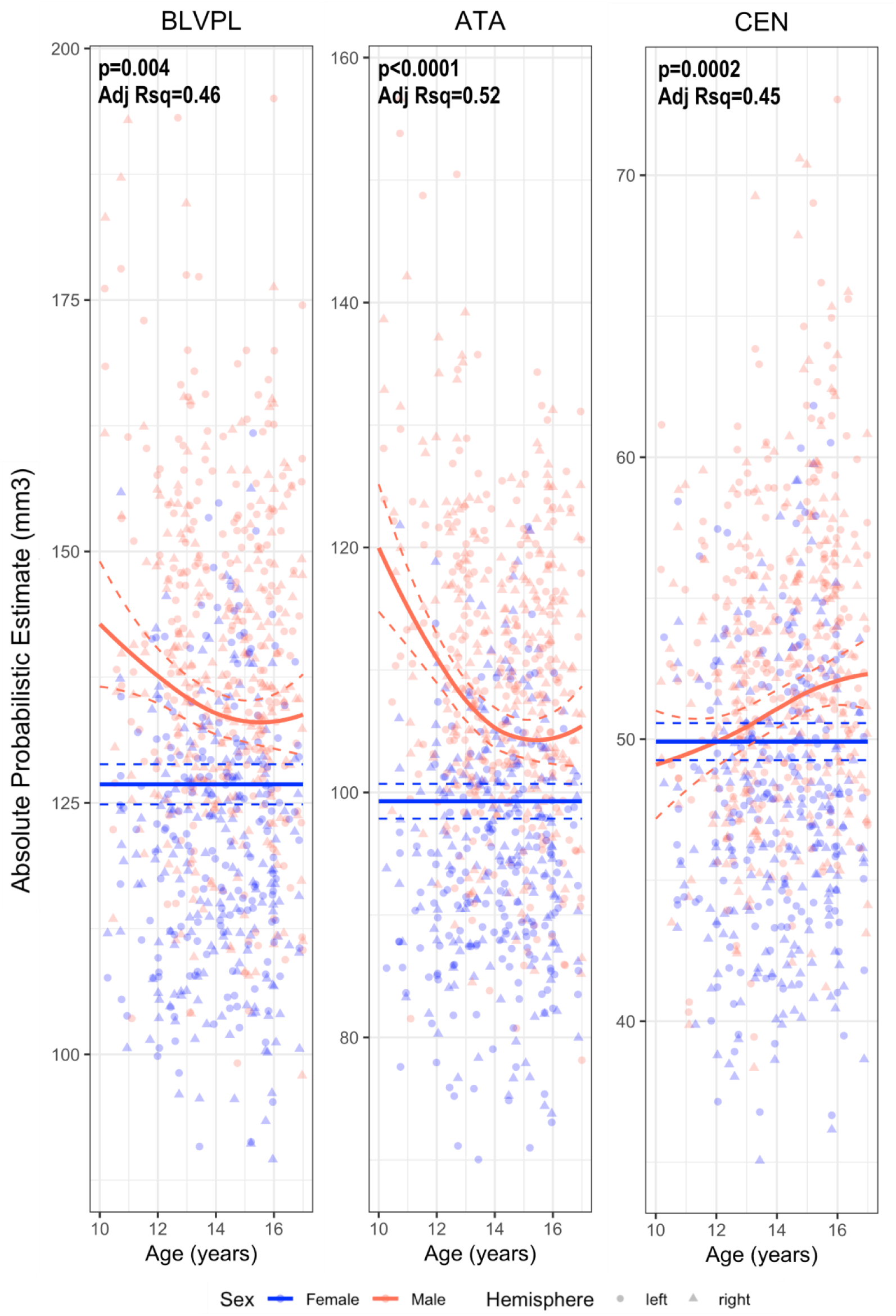
Sex differences in age associations with Absolute Probabilistic Estimate of the amygdala subregions. A) Basolateral ventral and paralaminar subdivision (BLVPL) and B) Central (CEN) and C) Amygdala transition area (ATA). Absolute Probabilistic Estimate plotted by age and sex (collapsed across hemispheres); solid lines reflect GAMM predicted fit estimates and dashed lines reflect 95% confidence intervals.

**Figure 5:**
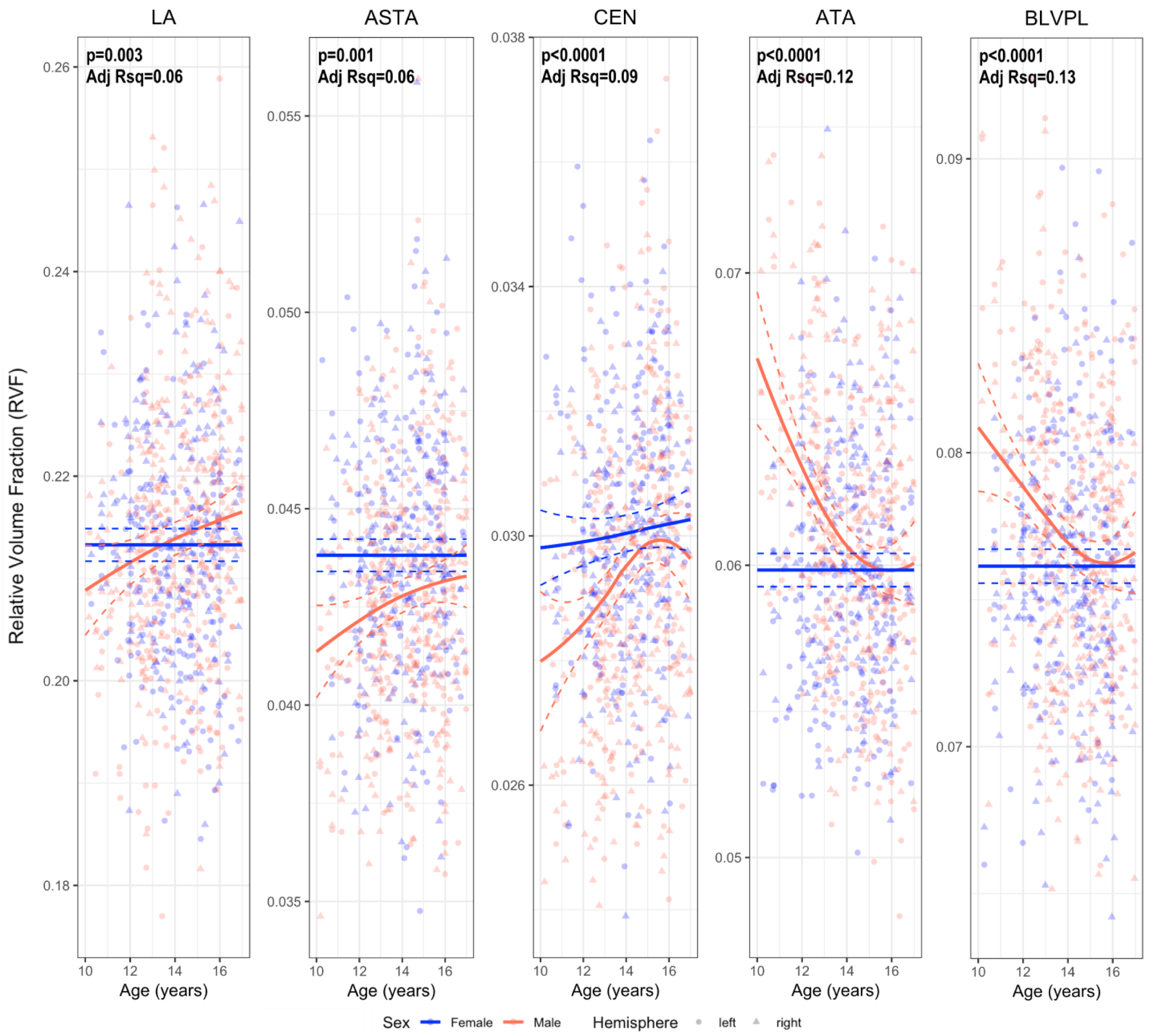
Sex differences in age associations with RVF of the amygdala subregions. A) Lateral nucleus (LA), B) Basolateral ventral and paralaminar subdivision (BLVPL) and C) Central (CEN) D) Amygdala transition area (ATA) and E) Amygdalostriatal transition area (ASTA). Relative Volume France (RVF) plotted by age and sex (collapsed across hemispheres); solid lines reflect GAMM predicted fit estimates and dashed lines reflect 95% confidence intervals.

### Amygdala subregions: Pubertal effects

GAMM model outputs for absolute probabilistic estimates (M4-M6) or *proportion* (i.e. RVF, M7-M9) for age, puberty, and age-by-puberty are presented for each sex separately. For females, no significant age, puberty, or age-by-puberty associations were seen for any of the 9 amygdala subregions (**Supplemental Tables 3-4**). In males, puberty and/or age-by-puberty effects were seen for the absolute volumes and proportion estimates of the BLVPL, CEN, and ATA (**Tables 4 and 5**). An age-by-pubertal interaction was seen for absolute BLVPL volumes (age-by-PDS: p=0.02; Adj R^2^: 0.27); though it did not pass multiple comparison correction. Moreover, although pubertal status was found to relate to the relative proportion of the BLVPL (M8: p’s ≤ 0.0056), the model with both age and pubertal status, as well as model comparison of the AIC and log-likelihood estimates, suggested age alone was the best predictor. For the CEN and ATA, both the absolute volumes as well as their relative proportions were found to significantly relate to pubertal status in males (M5 and M8: p’s ≤ 0.0056). Again, models including both age and pubertal status as well as AIC and log-likelihood suggested age was the best predictor. We also confirmed adding ICV as a covariate in proportion models (I.E. RVF) lead to similar results (**Supporting Tables 5 and 6**).

**Table 5.** GAMM amygdala subregion Absolute Probabilistic Estimate results for age, pubertal status, and age-by-pubertal status interaction for males.

| MALES |  |  |  |  |  |  |  |  |  |  |  |  |  |
| --- | --- | --- | --- | --- | --- | --- | --- | --- | --- | --- | --- | --- | --- |
| Total | Smooth Terms |  |  |  |  | Model Fit |  |  |  |  |  |  |  |
|  | Terms | edf | Ref.df | F | p-value | R2 | df | AIC | BIC | logLik | Test | L.Ratio | p-value |
| M4 | s(age) | 0.03 | 3.00 | 0.00 | 0.9176 | 0.3544 | 8 | 5615.69 | 5648.70 | -2799.84 | M4 vs. M6 | 1.83 | 0.7676 |
| M5 | s(pds) | 0.02 | 3.00 | 0.00 | 1.0000 | 0.3544 | 8 | 5615.69 | 5648.70 | -2799.84 | M5 vs. M6 | 1.83 | 0.7678 |
| M6 | ti(age) | 0.00 | 3.00 | 0.00 | 0.8394 | 0.3572 | 12 | 5621.86 | 5671.38 | -2798.93 |  |  |  |
|  | ti(pds) | 0.00 | 3.00 | 0.00 | 0.6553 |  |  |  |  |  |  |  |  |
|  | ti(age, pds) | 1.02 | 1.02 | 1.73 | 0.1847 |  |  |  |  |  |  |  |  |
| LA | Smooth Terms |  |  |  |  | Model Fit |  |  |  |  |  |  |  |
|  | Terms | edf | Ref.df | F | p-value | R2 | df | AIC | BIC | logLik | Test | L.Ratio | p-value |
| M4 | s(age) | 0.62 | 3.00 | 0.44 | 0.1398 | 0.2106 | 8 | 4539.26 | 4572.27 | -2261.63 | M4 vs. M6 | 0.11 | 0.9986 |
| M5 | s(pds) | 0.43 | 3.00 | 0.24 | 0.1958 | 0.2088 | 8 | 4539.52 | 4572.54 | -2261.76 | M5 vs. M6 | 0.37 | 0.9845 |
| M6 | ti(age) | 0.58 | 3.00 | 0.47 | 0.1141 | 0.2098 | 12 | 4547.15 | 4596.67 | -2261.57 |  |  |  |
|  | ti(pds) | 0.01 | 3.00 | 0.00 | 0.3013 |  |  |  |  |  |  |  |  |
|  | ti(age, pds) | 1.04 | 1.04 | 0.13 | 0.7324 |  |  |  |  |  |  |  |  |
| BLDI | Smooth Terms |  |  |  |  | Model Fit |  |  |  |  |  |  |  |
|  | Terms | edf | Ref.df | F | p-value | R2 | df | AIC | BIC | logLik | Test | L.Ratio | p-value |
| M4 | s(age) | 0.02 | 3.00 | 0.00 | 0.6726 | 0.2587 | 8 | 3930.65 | 3963.67 | -1957.33 | M4 vs. M6 | 1.96 | 0.7440 |
| M5 | s(pds) | 0.03 | 3.00 | 0.00 | 0.5643 | 0.2587 | 8 | 3930.65 | 3963.67 | -1957.33 | M5 vs. M6 | 1.95 | 0.7441 |
| M6 | ti(age) | 0.00 | 3.00 | 0.00 | 1.0000 | 0.2622 | 12 | 3936.70 | 3986.22 | -1956.35 |  |  |  |
|  | ti(pds) | 0.00 | 3.00 | 0.00 | 0.8437 |  |  |  |  |  |  |  |  |
|  | ti(age, pds) | 1.02 | 1.02 | 1.86 | 0.1691 |  |  |  |  |  |  |  |  |
| BLVPL | Smooth Terms |  |  |  |  | Model Fit |  |  |  |  |  |  |  |
|  | Terms | edf | Ref.df | F | p-value | R2 | df | AIC | BIC | logLik | Test | L.Ratio | p-value |
| M4 | s(age) | 1.64 | 3.00 | 2.55 | 0.0098 | 0.2559 | 8 | 3621.25 | 3654.27 | -1802.63 | M4 vs. M6 | 4.79 | 0.3095 |
| M5 | s(pds) | 0.74 | 3.00 | 0.69 | 0.0877 | 0.2405 | 8 | 3623.92 | 3656.94 | -1803.96 | M5 vs. M6 | 7.46 | 0.1136 |
| M6 | ti(age) | 0.06 | 3.00 | 0.03 | 0.2222 | 0.2697 | 12 | 3624.46 | 3673.99 | -1800.23 |  |  |  |
|  | ti(pds) | 0.02 | 3.00 | 0.00 | 0.3375 |  |  |  |  |  |  |  |  |
|  | ti(age, pds) | 3.31 | 3.31 | 3.15 | 0.0207 |  |  |  |  |  |  |  |  |
| BM | Smooth Terms |  |  |  |  | Model Fit |  |  |  |  |  |  |  |
|  | Terms | edf | Ref.df | F | p-value | R2 | df | AIC | BIC | logLik | Test | L.Ratio | p-value |
| M4 | s(age) | 0.02 | 3.00 | 0.00 | 0.6703 | 0.3370 | 8 | 3417.37 | 3450.39 | -1700.69 | M4 vs. M6 | 1.19 | 0.8792 |
| M5 | s(pds) | 0.02 | 3.00 | 0.00 | 0.6011 | 0.3370 | 8 | 3417.37 | 3450.39 | -1700.69 | M5 vs. M6 | 1.19 | 0.8792 |
| M6 | ti(age) | 0.00 | 3.00 | 0.00 | 0.8597 | 0.3382 | 12 | 3424.18 | 3473.70 | -1700.09 |  |  |  |
|  | ti(pds) | 0.00 | 3.00 | 0.00 | 0.8333 |  |  |  |  |  |  |  |  |
|  | ti(age, pds) | 1.01 | 1.01 | 1.15 | 0.2816 |  |  |  |  |  |  |  |  |
| CMN | Smooth Terms |  |  |  |  | Model Fit |  |  |  |  |  |  |  |
|  | Terms | edf | Ref.df | F | p-value | R2 | df | AIC | BIC | logLik | Test | L.Ratio | p-value |
| M4 | s(age) | 0.06 | 3.00 | 0.01 | 0.4113 | 0.3280 | 8 | 3709.99 | 3743.00 | -1846.99 | M4 vs. M6 | 2.97 | 0.5636 |
| M5 | s(pds) | 0.02 | 3.00 | 0.00 | 0.8086 | 0.3279 | 8 | 3709.98 | 3743.00 | -1846.99 | M5 vs. M6 | 2.96 | 0.5639 |
| M6 | ti(age) | 0.00 | 3.00 | 0.00 | 0.6853 | 0.3331 | 12 | 3715.02 | 3764.54 | -1845.51 |  |  |  |
|  | ti(pds) | 0.00 | 3.00 | 0.00 | 0.7963 |  |  |  |  |  |  |  |  |
|  | ti(age, pds) | 1.01 | 1.01 | 2.84 | 0.0893 |  |  |  |  |  |  |  |  |
| CEN | Smooth Terms |  |  |  |  | Model Fit |  |  |  |  |  |  |  |
|  | Terms | edf | Ref.df | F | p-value | R2 | df | AIC | BIC | logLik | Test | L.Ratio | p-value |
| M4 | s(age) | 1.54 | 3.00 | 4.38 | **0.0003** | 0.3246 | 8 | 2597.87 | 2630.88 | -1290.93 | M4 vs. M6 | 0.74 | 0.9466 |
| M5 | s(pds) | 1.07 | 3.00 | 2.55 | **0.0040** | 0.3121 | 8 | 2601.82 | 2634.83 | -1292.91 | M5 vs. M6 | 4.68 | 0.3212 |
| M6 | ti(age) | 2.12 | 3.00 | 5.82 | **0.0001** | 0.3337 | 12 | 2605.13 | 2654.65 | -1290.57 |  |  |  |
|  | ti(pds) | 0.00 | 3.00 | 0.00 | 0.8718 |  |  |  |  |  |  |  |  |
|  | ti(age, pds) | 1.01 | 1.01 | 1.24 | 0.2647 |  |  |  |  |  |  |  |  |
| AAA | Smooth Terms |  |  |  |  | Model Fit |  |  |  |  |  |  |  |
|  | Terms | edf | Ref.df | F | p-value | R2 | df | AIC | BIC | logLik | Test | L.Ratio | p-value |
| M4 | s(age) | 0.01 | 3.00 | 0.00 | 0.5225 | 0.2070 | 8 | 3050.85 | 3083.87 | -1517.43 | M4 vs. M6 | 1.61 | 0.8061 |
| M5 | s(pds) | 0.00 | 3.00 | 0.00 | 1.0000 | 0.2070 | 8 | 3050.85 | 3083.87 | -1517.43 | M5 vs. M6 | 1.61 | 0.8063 |
| M6 | ti(age) | 0.00 | 3.00 | 0.00 | 1.0000 | 0.2138 | 12 | 3057.24 | 3106.76 | -1516.62 |  |  |  |
|  | ti(pds) | 0.00 | 3.00 | 0.00 | 0.7833 |  |  |  |  |  |  |  |  |
|  | ti(age, pds) | 2.37 | 2.37 | 2.15 | 0.2846 |  |  |  |  |  |  |  |  |
| ATA | Smooth Terms |  |  |  |  | Model Fit |  |  |  |  |  |  |  |
|  | Terms | edf | Ref.df | F | p-value | R2 | df | AIC | BIC | logLik | Test | L.Ratio | p-value |
| M4 | s(age) | 2.32 | 3.00 | 11.74 | **1.57E-08** | 0.3339 | 8 | 3347.79 | 3380.80 | -1665.89 | M4 vs. M6 | 1.13 | 0.8892 |
| M5 | s(pds) | 1.26 | 3.00 | 3.70 | **0.0008** | 0.2828 | 8 | 3365.61 | 3398.62 | -1674.80 | M5 vs. M6 | 18.95 | **0.0008** |
| M6 | ti(age) | 1.46 | 3.00 | 4.84 | **0.0001** | 0.3294 | 12 | 3354.66 | 3404.18 | -1665.33 |  |  |  |
|  | ti(pds) | 0.01 | 3.00 | 0.00 | 0.4392 |  |  |  |  |  |  |  |  |
|  | ti(age, pds) | 1.13 | 1.13 | 2.52 | 0.0904 |  |  |  |  |  |  |  |  |
| ASTA | Smooth Terms |  |  |  |  | Model Fit |  |  |  |  |  |  |  |
|  | Terms | edf | Ref.df | F | p-value | R2 | df | AIC | BIC | logLik | Test | L.Ratio | p-value |
| M4 | s(age) | 0.92 | 3.00 | 1.15 | 0.0401 | 0.2312 | 8 | 2989.47 | 3022.48 | -1486.73 | M4 vs. M6 | 0.01 | 1.0000 |
| M5 | s(pds) | 0.62 | 3.00 | 0.47 | 0.1292 | 0.2255 | 8 | 2990.84 | 3023.85 | -1487.42 | M5 vs. M6 | 1.36 | 0.8518 |
| M6 | ti(age) | 0.80 | 3.00 | 1.03 | 0.0425 | 0.2295 | 12 | 2997.48 | 3047.00 | -1486.74 |  |  |  |
|  | ti(pds) | 0.01 | 3.00 | 0.00 | 0.6686 |  |  |  |  |  |  |  |  |
|  | ti(age, pds) | 1.02 | 1.02 | 0.01 | 0.9082 |  |  |  |  |  |  |  |  |

Notes: In each model, the smooth terms, the estimated degree of freedom (edf), reference degree of freedom (Ref.df), F-score, p-value, and Adjusted R<sup>2</sup> for each model is shown; p-value < 0.0056 bolded (Bonferroni corrected). Between model comparisons include the df, AIC, log-likelihood ratio (L Ratio) and p-values < 0.05 bolded. **Table 6. GAMM amygdala subregion Relative Volume Fraction (RVF) results for age, pubertal status, and age-** **by-pubertal status interaction for males**
Notes: In each model, the smooth terms, the estimated degree of freedom (edf), reference degree of freedom (Ref.df), F-score, p-value, and Adjusted R<sup>2</sup> for each model is shown; p-value < 0.0056 bolded (Bonferroni corrected). Between model comparisons include the df, AIC, log-likelihood ratio (L Ratio) and p-values < 0.05 bolded.

**Table 6.** GAMM amygdala subregion Relative Volume Fraction (RVF) results for age, pubertal status, and age-by-pubertal status interaction for males.

| MALES |  |  |  |  |  |  |  |  |  |  |  |  |  |
| --- | --- | --- | --- | --- | --- | --- | --- | --- | --- | --- | --- | --- | --- |
| LA | Smooth Terms |  |  |  |  | Model Fit |  |  |  |  |  |  |  |
|  | Terms | edf | Ref.df | F | p-value | R2 | df | AIC | BIC | logLik | Test | L.Ratio | p-value |
| M7 | s(age) | 1.22 | 3.00 | 2.32 | 0.0069 | 0.0745 | 7 | -2779.68 | -2750.79 | 1396.84 | M7 vs. M9 | 0.29 | 0.9903 |
| M8 | s(pds) | 0.90 | 3.00 | 1.30 | 0.0300 | 0.0650 | 7 | -2777.51 | -2748.62 | 1395.76 | M8 vs. M9 | 2.46 | 0.6514 |
| M9 | ti(age) | 0.95 | 3.00 | 1.77 | 0.0132 | 0.0722 | 11 | -2771.97 | -2726.58 | 1396.99 |  |  |  |
|  | ti(pds) | 0.00 | 3.00 | 0.00 | 0.6037 |  |  |  |  |  |  |  |  |
|  | ti(age, pds) | 1.00 | 1.00 | 0.34 | 0.5605 |  |  |  |  |  |  |  |  |
| BLDI | Smooth Terms |  |  |  |  | Model Fit |  |  |  |  |  |  |  |
|  | Terms | edf | Ref.df | F | p-value | R2 | df | AIC | BIC | logLik | Test | L.Ratio | p-value |
| M7 | s(age) | 0.49 | 3.00 | 0.25 | 0.2163 | 0.1388 | 7 | -3876.94 | -3848.05 | 1945.47 | M7 vs. M9 | 1.66 | 0.7988 |
| M8 | s(pds) | 0.80 | 3.00 | 0.95 | 0.0526 | 0.1451 | 7 | -3878.31 | -3849.42 | 1946.15 | M8 vs. M9 | 0.28 | 0.9909 |
| M9 | ti(age) | 0.00 | 3.00 | 0.00 | 0.8181 | 0.1436 | 11 | -3870.59 | -3825.20 | 1946.30 |  |  |  |
|  | ti(pds) | 0.69 | 3.00 | 0.71 | 0.0757 |  |  |  |  |  |  |  |  |
|  | ti(age, pds) | 1.01 | 1.01 | 0.29 | 0.5876 |  |  |  |  |  |  |  |  |
| BLVPL | Smooth Terms |  |  |  |  | Model Fit |  |  |  |  |  |  |  |
|  | Terms | edf | Ref.df | F | p-value | R2 | df | AIC | BIC | logLik | Test | L.Ratio | p-value |
| M7 | s(age) | 2.32 | 3.00 | 10.61 | 8.00E-08 | 0.1392 | 7 | -3629.03 | -3600.14 | 1821.51 | M7 vs. M9 | 3.14 | 0.5345 |
| M8 | s(pds) | 1.11 | 3.00 | 2.95 | 0.0021 | 0.1062 | 7 | -3677.58 | -3648.69 | 1845.79 | M8 vs. M9 | 45.41 | 3.26E-09 |
| M9 | ti(age) | 1.98 | 3.00 | 6.29 | 7.09E-06 | 0.1718 | 11 | -3624.17 | -3578.77 | 1823.08 |  |  |  |
|  | ti(pds) | 0.01 | 3.00 | 0.00 | 0.2791 |  |  |  |  |  |  |  |  |
|  | ti(age, pds) | 5.63 | 5.63 | 2.39 | 0.1672 |  |  |  |  |  |  |  |  |
| BM | Smooth Terms |  |  |  |  | Model Fit |  |  |  |  |  |  |  |
|  | Terms | edf | Ref.df | F | p-value | R2 | df | AIC | BIC | logLik | Test | L.Ratio | p-value |
| M7 | s(age) | 0.00 | 3.00 | 0.00 | 0.4152 | 0.0394 | 7 | -3863.77 | -3834.89 | 1938.89 | M7 vs. M9 | 0.95 | 0.9175 |
| M8 | s(pds) | 0.30 | 3.00 | 0.13 | 0.2510 | 0.0402 | 7 | -3863.82 | -3834.93 | 1938.91 | M8 vs. M9 | 0.90 | 0.9242 |
| M9 | ti(age) | 0.00 | 3.00 | 0.00 | 0.6623 | 0.0446 | 11 | -3856.72 | -3811.33 | 1939.36 |  |  |  |
|  | ti(pds) | 0.01 | 3.00 | 0.00 | 0.4209 |  |  |  |  |  |  |  |  |
|  | ti(age, pds) | 2.40 | 2.40 | 0.71 | 0.4121 |  |  |  |  |  |  |  |  |
| CMN | Smooth Terms |  |  |  |  | Model Fit |  |  |  |  |  |  |  |
|  | Terms | edf | Ref.df | F | p-value | R2 | df | AIC | BIC | logLik | Test | L.Ratio | p-value |
| M7 | s(age) | 0.09 | 3.00 | 0.03 | 0.3300 | 0.0208 | 7 | -3502.36 | -3473.47 | 1758.18 | M7 vs. M9 | 0.29 | 0.9902 |
| M8 | s(pds) | 0.02 | 3.00 | 0.00 | 0.7752 | 0.0204 | 7 | -3502.35 | -3473.46 | 1758.18 | M8 vs. M9 | 0.30 | 0.9896 |
| M9 | ti(age) | 0.00 | 3.00 | 0.00 | 0.3768 | 0.0195 | 11 | -3494.65 | -3449.26 | 1758.33 |  |  |  |
|  | ti(pds) | 0.00 | 3.00 | 0.00 | 0.9123 |  |  |  |  |  |  |  |  |
|  | ti(age, pds) | 1.05 | 1.05 | 0.26 | 0.6056 |  |  |  |  |  |  |  |  |
| CEN | Smooth Terms |  |  |  |  | Model Fit |  |  |  |  |  |  |  |
|  | Terms | edf | Ref.df | F | p-value | R2 | df | AIC | BIC | logLik | Test | L.Ratio | p-value |
| M7 | s(age) | 2.53 | 3.00 | 11.10 | 8.04E-08 | 0.0904 | 7 | -4274.82 | -4245.93 | 2144.41 | M7 vs. M9 | 0.65 | 0.9573 |
| M8 | s(pds) | 1.10 | 3.00 | 2.94 | 0.0020 | 0.0526 | 7 | -4323.70 | -4294.82 | 2168.85 | M8 vs. M9 | 48.23 | 8.44E-10 |
| M9 | ti(age) | 2.51 | 3.00 | 10.09 | 1.72E-07 | 0.0920 | 11 | -4267.47 | -4222.08 | 2144.74 |  |  |  |
|  | ti(pds) | 0.00 | 3.00 | 0.00 | 0.7837 |  |  |  |  |  |  |  |  |
|  | ti(age, pds) | 1.10 | 1.10 | 0.70 | 0.4312 |  |  |  |  |  |  |  |  |
| AAA | Smooth Terms |  |  |  |  | Model Fit |  |  |  |  |  |  |  |
|  | Terms | edf | Ref.df | F | p-value | R2 | df | AIC | BIC | logLik | Test | L.Ratio | p-value |
| M7 | s(age) | 0.01 | 3.00 | 0.00 | 0.5058 | -0.0058 | 7 | -4121.01 | -4092.13 | 2067.51 | M7 vs. M9 | 1.46 | 0.8331 |
| M8 | s(pds) | 0.01 | 3.00 | 0.00 | 0.9963 | -0.0058 | 7 | -4121.01 | -4092.12 | 2067.50 | M8 vs. M9 | 1.47 | 0.8321 |
| M9 | ti(age) | 0.02 | 3.00 | 0.00 | 0.4720 | 0.0077 | 11 | -4114.48 | -4069.08 | 2068.24 |  |  |  |
|  | ti(pds) | 0.01 | 3.00 | 0.00 | 0.9471 |  |  |  |  |  |  |  |  |
|  | ti(age, pds) | 3.56 | 3.56 | 1.65 | 0.2284 |  |  |  |  |  |  |  |  |
| ATA | Smooth Terms |  |  |  |  | Model Fit |  |  |  |  |  |  |  |
|  | Terms | edf | Ref.df | F | p-value | R2 | df | AIC | BIC | logLik | Test | L.Ratio | p-value |
| M7 | s(age) | 2.47 | 3.00 | 22.82 | 1.80E-15 | 0.1556 | 7 | -3672.81 | -3643.93 | 1843.41 | M7 vs. M9 | 59.58 | 3.56E-12 |
| M8 | s(pds) | 1.31 | 3.00 | 5.59 | 3.35E-05 | 0.0819 | 7 | -3711.02 | -3682.13 | 1862.51 | M8 vs. M9 | 21.38 | 0.0003 |
| M9 | ti(age) | 2.16 | 3.00 | 10.72 | 2.36E-08 | 0.1546 | 11 | -3724.39 | -3679.00 | 1873.20 |  |  |  |
|  | ti(pds) | 0.00 | 3.00 | 0.00 | 0.5055 |  |  |  |  |  |  |  |  |
|  | ti(age, pds) | 1.01 | 1.01 | 0.03 | 0.8626 |  |  |  |  |  |  |  |  |
| ASTA | Smooth Terms |  |  |  |  | Model Fit |  |  |  |  |  |  |  |
|  | Terms | edf | Ref.df | F | p-value | R2 | df | AIC | BIC | logLik | Test | L.Ratio | p-value |
| M7 | s(age) | 1.62 | 3.00 | 4.10 | 0.0006 | 0.0328 | 7 | -3980.89 | -3952.00 | 1997.44 | M7 vs. M9 | 2.69 | 0.6107 |
| M8 | s(pds) | 0.76 | 3.00 | 0.80 | 0.0695 | 0.0153 | 7 | -4034.94 | -4006.05 | 2024.47 | M8 vs. M9 | 51.36 | 1.88E-10 |
| M9 | ti(age) | 0.75 | 3.00 | 0.91 | 0.0427 | 0.0409 | 11 | -3975.58 | -3930.18 | 1998.79 |  |  |  |
|  | ti(pds) | 0.02 | 3.00 | 0.00 | 0.3549 |  |  |  |  |  |  |  |  |
|  | ti(age, pds) | 3.02 | 3.02 | 2.08 | 0.0904 |  |  |  |  |  |  |  |  |

Notes: In each model, the smooth terms, the estimated degree of freedom (edf), reference degree of freedom (Ref.df), F-score, p-value, and Adjusted R<sup>2</sup> for each model is shown; p-value < 0.0056 bolded (Bonferroni corrected). Between model comparisons include the df, AIC, log-likelihood ratio (L Ratio) and p-values < 0.05 bolded.

## Discussion

The current cross-sectional study provides the first glimpse at associations between amygdala subregion volumes and apportionment across adolescence. While previous studies have examined total amygdala volumes across childhood and adolescence (M. M. Herting et al., 2018; L. M. Wierenga et al., 2018), the current study highlights the utility of the CIT168 to define 9 amygdala subregions in a large sample of adolescents and suggests that within the amygdala distinct subregions may evolve across the adolescent period in a sex specific fashion. Using the newly derived *in vivo* CIT168 atlas to examine subregional volumes and proportional estimates, we found larger CEN, but smaller BLVPL and ATA subregion volumes with age in males, but not females. In addition, as a function of age-related differences in subregional volumes, the relative proportional findings suggest 17 year-old males have proportionately larger LA, CEN, and ASTA, but smaller BLVPL and ATA subregions, compared to 10 year-old males. Interestingly, the age associations in the apportionment of major subregions in males results in greater similarity of amygdala apportionment to females by age 17 years as compared to earlier on at age 10.

Although cross-sectional, our findings lend support for the hypothesis that subregions within the amygdala may have unique development patterns in males during the adolescent years. While future longitudinal MRI studies are needed to confirm potential changes in subregional volumes across development *in vivo*, our findings in males are supported by the recent histological study showing that postnatal neuron numbers change in distinct nuclei, including the central, lateral, and basal nuclei, from childhood to adulthood (Avino et al., 2018). In that study, however, a sex-specific effect was not examined, as the sample included a wide age range (n=24, 2 to 48 years) with very few females in the neurotypical subgroup (n = 5) (Avino et al., 2018). Beyond nucleus-specific changes in neuron number, postnatal immunohistochemistry studies have also found a difference in immature and mature neuron concentrations among amygdala nuclei, including the lateral, central, basal, and paralaminar nucleus (Avino et al., 2018). A higher concentration of immature neurons has been reported in the paralaminar nucleus (part of the BLVPL subregion in the current study) as compared to other amygdala nuclei (Avino et al., 2018). Moreover, the number of immature neurons in the paralaminar nucleus decreases over time, whereas the mature neuron numbers of the surrounding regions continue to increase in childhood and adolescence. These data have led to the hypothesis that gradual maturation and migration of paralaminar immature neurons may contribute to the mature neuron number within the paralaminar, and/or be the source of increases in neuron number seen in other nuclei over development (Avino et al., 2018; deCampo & Fudge, 2012). If this hypothesis proves to be correct, migration and maturation of immature neurons may contribute to the size as well as the re-configuration and/or refinement of the amygdala subregions and their subsequent connectivity with the cerebral cortex across adolescence. While MRI cannot assess neuron number, more research is needed to determine if age associations with the absolute volumes of the BLVPL, CEN, and ATA as well as age related differences in the relative proportion of these amygdala subregions, along with the LA and ASTA, in males may be suggestive of distinct nuclei maturation and migration patterns in amygdala development. Combining postmortem histology and MRI segmentation approaches in developing samples is necessary to further decipher if these age and sex-specific patterns occur across development.

Furthermore, cytoarchitectural findings suggest the BLVPL subregion of the amygdala receives afferents from both the lateral nucleus (LA) and the hippocampus (Pitkanen & Amaral, 1998). Efferents of the medial paralaminar nucleus gradually merge with the periamygdaloid cortex, corresponding to the amygdaloid transition areas (ATA) in this study, which in turn projects to the hippocampus. Moreover, the lateral nucleus (LA) receives sensory information, allowing the basolateral complex to process the information, and then send this information out of the amygdala via the central nucleus (CEN) (McDonald & Jackson, 1987; Sah, Faber, Lopez De Armentia, & Power, 2003). The CIT168 ATA region encapsulates the periamygdaloid cortex, as well as these amygdalocortical and amygdolohippocampal transition areas. Hippocampal input to the amygdala is important for contextual fear learning (Phillips & LeDoux, 1992), and given the convergence between sensory input from the LA, as well as bidirectional connectivity with the hippocampus, it has been proposed that the paralaminar and periamygdaloid cortex of the amygdala may be involved in contextual learning (deCampo & Fudge, 2013). It remains to be elucidated how larger LA, ASTA, and CEN, but smaller BLVPL and ATA apportionment in males may map onto function. However, amygdala nuclei size and relative composition may be an additional MRI feature to explore in hopes of clarifying our understanding of amygdala structural and functional development. It may also prove useful in studying known sex differences in emotion-related behavior, brain function, and prevalence in mental health disorders that typically emerge during this time. For example, meta-analysis of 166 studies found a small, yet consistent, sex difference in positive and negative emotional expression that begins to diverge in the beginning of childhood and into adolescence (Chaplin & Aldao, 2013). Similarly, fMRI studies have reported greater brain activity in cortical regions, including visual and parietal regions, in male versus female adolescents during emotional functional MRI tasks (Cservenka, Stroup, Etkin, & Nagel, 2015). Resting-state fMRI studies implementing *ex vivo* atlases to define basolateral, superficial, and centromedial subregions, have also found age and sex-specific differences in amygdala functional connectivity patterns. Age and region-specific patterns were seen between the amygdala and medial prefrontal cortex, with connectivity becoming apparent at age 10 and continuing to strengthen across adolescence (Gabard-Durnam et al., 2014). In a separate study, the superficial amygdala resting-state patterns were found to be more mature in female adolescents, but basolateral amygdala connectivity patterns were more mature in male adolescents (Alarcon et al., 2015). Future studies are warranted to determine if the absolute and/or relative differences in the volumes of primary input (LA) and output (CEN) subregions, as well as subregions involved in contextual and emotional learning (BLVPL, ATA) in males, may relate to differences in emotional expression, greater cortical activation to emotional stimuli and/or stronger basolateral functional connectivity in males versus females during adolescence. Beyond the possible functional implications of nuclei apportionment, implementation of the CIT168 atlas to construct ROIs for other MRI modalities, including resting-state fMRI, task-based fMRI, and diffusion, may also assist in gaining greater specificity of how different amygdala nuclei functionally and structurally develop.

While this is the first study to examine amygdala subregional volume composition in adolescents, the current study has both strengths and limitations. Other amygdala segmentation approaches are derived from post-mortem samples that are largely based on smaller samples of older male brains (Amunts et al., 2005; Saygin et al., 2017), which not only fail to capture possible developmental changes but may also be confounded by factors that influence tissue quality (Stan et al., 2006). The CIT168 atlas mitigates some of these concerns by using the high-resolution (700 micrometer) *in vivo* Human Connectome Project data from young adults (ages 22-35 years) and by using probabilistic delineations to encode partial volume uncertainty in amygdala subregion boundaries. While the CIT168 probabilistic amygdala atlas has several advantages over other currently available approaches, challenges still exist when registering templates derived from *in vivo* adult human data to anatomic data acquired in developing human children on different scanners, at different field strengths, and with differing imaging parameters. Amygdala nuclei remain extremely difficult to discern by eye in typical 3T whole brain T1- or T2-weighted images, including those in the current study. While we have used conservative exclusion criteria for minimum CNR to improve the robustness of our volume estimates, replication studies at higher field strengths (e.g. 7 Tesla) with better individual-level amygdala CNR would be extremely valuable. A few additional limitations should also be noted. First, the current study is cross-sectional and therefore cannot attest to development of the amygdala over time in the same individuals. Future longitudinal studies are necessary and warranted to further elucidate potential maturational patterns of amygdala subregion volumes and apportionment in each sex and as individuals undergo pubertal maturation. From this cross-sectional analysis, our hypothesis that pubertal development would relate to amygdala composition during adolescence was not well supported. While physical characteristics of pubertal maturation did relate to BLVPL, CEN, and ATA, our current results suggest that age alone best accounts for individual differences in amygdala nuclei volume composition in males. Moreover, neither age nor pubertal status related to any of the nuclei examined in females. It is possible the lack of associations is due to our study sample. Although pubertal development scores were on average similar between the sexes in our sample (**Table 1**), there were fewer females that fell within the pre-pubertal and early pubertal range as compared to males in this age range of 10 to 17 years. While this is to be expected given the known sex difference in pubertal onset, with girls showing physical signs of maturation ∼1-2 years prior to males (Dorn, 2006), more research is needed in younger females in order to assess if similar patterns of amygdala maturation do occur at slightly younger ages in females. Furthermore, as a focus of future research, it would also be helpful to utilize other markers that may be more accurate for capturing both puberty in children, such as pubertal hormone levels.

To summarize, we show the adolescent amygdala can be segmented into 9 subregions using the newly developed CIT168 atlas and that the relative composition of these amygdala subregions may continue to restructure in a sex-specific fashion during the adolescent window of development. By using this approach in conjunction with considering how the amygdala nuclei composition may continue to develop, future studies may be able to further explore how the amygdaloid complex may interact with distinct cortical regions, such as the prefrontal cortex, in order to modulate each other’s development and social and emotional behaviors that continue to mature during this critical period in development (Andersen & Teicher, 2008; Tottenham & Gabard-Durnam, 2017). Our approach provides a first step towards a more rigorous exploration of functional and structural connectivity development within the heterogeneous amygdala complex across adolescence.

## Acknowledgements

The research above was supported by the following grants, R01 AA017664 (PI: Nagel), R21 MH099618 (PI: Nagel), R03 HD090308 (PI: Herting), K01 MH108761 (PI: Herting), and NIMH P50 MH094258 #8198 (PI: Tyszka). We also thank the families who contributed their time and participated in the above study.

## Conflict of Interest

The authors have no conflict of interests to declare.

## Data Availability Statement

Data available on request from the authors.

## Notes

### Competing Interest Statement

The authors have declared no competing interest.

## References

1. Alarcon, G., Cservenka, A., Rudolph, M. D., Fair, D. A., & Nagel, B. J. (2015). Developmental sex differences in resting state functional connectivity of amygdala sub-regions. Neuroimage, 115, 235–244. doi: 10.1016/j.neuroimage.2015.04.013

2. Amaral, D. G., & Price, J. L. (1984). Amygdalo-cortical projections in the monkey (Macaca fascicularis). J Comp Neurol, 230(4), 465–496. doi: 10.1002/cne.902300402

3. Amunts, K., Kedo, O., Kindler, M., Pieperhoff, P., Mohlberg, H., Shah, N. J., … Zilles, K. (2005). Cytoarchitectonic mapping of the human amygdala, hippocampal region and entorhinal cortex: intersubject variability and probability maps. Anat Embryol (Berl), 210(5-6), 343–352. doi: 10.1007/s00429-005-0025-5

4. Andersen, S. L., & Teicher, M. H. (2008). Stress, sensitive periods and maturational events in adolescent depression. Trends Neurosci, 31(4), 183–191. doi: 10.1016/j.tins.2008.01.004

5. Avants, B., Anderson, C., Grossman, M., & Gee, J. C. (2007). Spatiotemporal normalization for longitudinal analysis of gray matter atrophy in frontotemporal dementia. Med Image Comput Comput Assist Interv, 10(Pt 2), 303–310.

6. Avants, B. B., Tustison, N. J., Song, G., Cook, P. A., Klein, A., & Gee, J. C. (2011). A reproducible evaluation of ANTs similarity metric performance in brain image registration. Neuroimage, 54(3), 2033–2044. doi: 10.1016/j.neuroimage.2010.09.025

7. Avino, T. A., Barger, N., Vargas, M. V., Carlson, E. L., Amaral, D. G., Bauman, M. D., & Schumann, C. M. (2018). Neuron numbers increase in the human amygdala from birth to adulthood, but not in autism. Proc Natl Acad Sci U S A, 115(14), 3710–3715. doi: 10.1073/pnas.1801912115

8. Backhausen, L. L., Herting, M. M., Buse, J., Roessner, V., Smolka, M. N., & Vetter, N. C. (2016). Quality Control of Structural MRI Images Applied Using FreeSurfer-A Hands-On Workflow to Rate Motion Artifacts. Front Neurosci, 10, 558. doi: 10.3389/fnins.2016.00558

9. Barbas, H., & De Olmos, J. (1990). Projections from the amygdala to basoventral and mediodorsal prefrontal regions in the rhesus monkey. J Comp Neurol, 300(4), 549–571. doi: 10.1002/cne.903000409

10. Baxter, M. G., & Murray, E. A. (2002). The amygdala and reward. Nat Rev Neurosci, 3(7), 563–573. doi: 10.1038/nrn875

11. Berenbaum, S. A., Beltz, A. M., & Corley, R. (2015). The importance of puberty for adolescent development: conceptualization and measurement. Adv Child Dev Behav, 48, 53–92. doi: 10.1016/bs.acdb.2014.11.002

12. Bernier, P. J., Bedard, A., Vinet, J., Levesque, M., & Parent, A. (2002). Newly generated neurons in the amygdala and adjoining cortex of adult primates. Proc Natl Acad Sci U S A, 99(17), 11464–11469. doi: 10.1073/pnas.172403999

13. Bonferroni, C. E. (1936). Teoria statistica delle classi e calcolo delle probabilita. Pubblicazioni del R Istituto Superiore di Sceinze Economiche e Commerciali di Firenze, 8, 3–62.

14. Bramen, J. E., Hranilovich, J. A., Dahl, R. E., Forbes, E. E., Chen, J., Toga, A. W., … Sowell, E. R. (2011). Puberty influences medial temporal lobe and cortical gray matter maturation differently in boys than girls matched for sexual maturity. Cereb Cortex, 21(3), 636–646. doi: 10.1093/cercor/bhq137

15. Brito, N. H., & Noble, K. G. (2014). Socioeconomic status and structural brain development. Front Neurosci, 8, 276. doi: 10.3389/fnins.2014.00276

16. Bzdok, D., Laird, A. R., Zilles, K., Fox, P. T., & Eickhoff, S. B. (2013). An investigation of the structural, connectional, and functional subspecialization in the human amygdala. Hum Brain Mapp, 34(12), 3247–3266. doi: 10.1002/hbm.22138

17. Chaplin, T. M., & Aldao, A. (2013). Gender differences in emotion expression in children: a meta-analytic review. Psychol Bull, 139(4), 735–765. doi: 10.1037/a0030737

18. Cservenka, A., Stroup, M. L., Etkin, A., & Nagel, B. J. (2015). The effects of age, sex, and hormones on emotional conflict-related brain response during adolescence. Brain Cogn, 99, 135–150. doi: 10.1016/j.bandc.2015.06.002

19. Cunningham, M. G., Bhattacharyya, S., & Benes, F. M. (2002). Amygdalo-cortical sprouting continues into early adulthood: implications for the development of normal and abnormal function during adolescence. J Comp Neurol, 453(2), 116–130. doi: 10.1002/cne.10376

20. deCampo, D. M., & Fudge, J. L. (2012). Where and what is the paralaminar nucleus? A review on a unique and frequently overlooked area of the primate amygdala. Neurosci Biobehav Rev, 36(1), 520–535. doi: 10.1016/j.neubiorev.2011.08.007

21. deCampo, D. M., & Fudge, J. L. (2013). Amygdala projections to the lateral bed nucleus of the stria terminalis in the macaque: comparison with ventral striatal afferents. J Comp Neurol, 521(14), 3191–3216. doi: 10.1002/cne.23340

22. Dorn, L. D. (2006). Measuring puberty. J Adolesc Health, 39(5), 625–626. doi: 10.1016/j.jadohealth.2006.05.014

23. Gabard-Durnam, L. J., Flannery, J., Goff, B., Gee, D. G., Humphreys, K. L., Telzer, E., … Tottenham, N. (2014). The development of human amygdala functional connectivity at rest from 4 to 23 years: a cross-sectional study. Neuroimage, 95, 193–207. doi: 10.1016/j.neuroimage.2014.03.038

24. Ghashghaei, H. T., & Barbas, H. (2002). Pathways for emotion: interactions of prefrontal and anterior temporal pathways in the amygdala of the rhesus monkey. Neuroscience, 115(4), 1261-1279. Retrieved from https://ac.els-cdn.com/S0306452202004463/1-s2.0-S0306452202004463-main.pdf?_tid=e8ac9f70-e2fc-4dc8-80ed-3a5c9f4c6bca&acdnat=1530044158_16bdb26c73bcc4a16b517bac63de410e

25. Giedd, J. N., Vaituzis, A. C., Hamburger, S. D., Lange, N., Rajapakse, J. C., Kaysen, D., … Rapoport, J. L. (1996). Quantitative MRI of the temporal lobe, amygdala, and hippocampus in normal human development: ages 4-18 years. J Comp Neurol, 366(2), 223–230. doi: 10.1002/(SICI)1096-9861(19960304)366:2<223::AID-CNE3>3.0.CO;2-7

26. Goddings, A. L., Mills, K. L., Clasen, L. S., Giedd, J. N., Viner, R. M., & Blakemore, S. J. (2014). The influence of puberty on subcortical brain development. Neuroimage, 88, 242–251. doi: 10.1016/j.neuroimage.2013.09.073

27. Hariri, A. R., Tessitore, A., Mattay, V. S., Fera, F., & Weinberger, D. R. (2002). The amygdala response to emotional stimuli: a comparison of faces and scenes. Neuroimage, 17(1), 317-323. Retrieved from https://ac.els-cdn.com/S1053811902911791/1-s2.0-S1053811902911791-main.pdf?_tid=d31ead47-dc11-4894-ba30-0010e3ac1968&acdnat=1531161816_b9d533b625b7f466bda500ee78b5cedb

28. Herting, M. M., Azad, A., Kim, R., Tyszka, J. M., Geffner, M. E., & Kim, M. S. (2020). Brain Differences in the Prefrontal Cortex, Amygdala, and Hippocampus in Youth with Congenital Adrenal Hyperplasia. The Journal of Clinical Endocrinology & Metabolism.

29. Herting, M. M., Gautam, P., Spielberg, J. M., Kan, E., Dahl, R. E., & Sowell, E. R. (2014). The role of testosterone and estradiol in brain volume changes across adolescence: a longitudinal structural MRI study. Hum Brain Mapp, 35(11), 5633–5645. doi: 10.1002/hbm.22575

30. Herting, M. M., Johnson, C., Mills, K. L., Vijayakumar, N., Dennison, M., Liu, C., … Tamnes, C. K. (2018). Development of subcortical volumes across adolescence in males and females: A multisample study of longitudinal changes. Neuroimage, 172, 194–205. doi: 10.1016/j.neuroimage.2018.01.020

31. Hollingshead, A. A. (1975). Four-factor index of social status. Unpublished manuscript, Yale University, New Haven, CT.

32. Janak, P. H., & Tye, K. M. (2015). From circuits to behaviour in the amygdala. Nature, 517(7534), 284–292. doi: 10.1038/nature14188

33. Jenkinson, M., Beckmann, C. F., Behrens, T. E., Woolrich, M. W., & Smith, S. M. (2012). FSL. Neuroimage, 62(2), 782–790. doi: 10.1016/j.neuroimage.2011.09.015

34. Killcross, S., Robbins, T. W., & Everitt, B. J. (1997). Different types of fear-conditioned behaviour mediated by separate nuclei within amygdala. Nature, 388(6640), 377–380. doi: 10.1038/41097

35. Krettek, J. E., & Price, J. L. (1978). A description of the amygdaloid complex in the rat and cat with observations on intra-amygdaloid axonal connections. J Comp Neurol, 178(2), 255–280. doi: 10.1002/cne.901780205

36. McDonald, A. J., & Jackson, T. R. (1987). Amygdaloid connections with posterior insular and temporal cortical areas in the rat. J Comp Neurol, 262(1), 59–77. doi: 10.1002/cne.902620106

37. Meyer-Lindenberg, A., Hariri, A. R., Munoz, K. E., Mervis, C. B., Mattay, V. S., Morris, C. A., & Berman, K. F. (2005). Neural correlates of genetically abnormal social cognition in Williams syndrome. Nat Neurosci, 8(8), 991–993. doi: 10.1038/nn1494

38. Morales, A. M., Jones, S. A., Ehlers, A., Lavine, J. B., & Nagel, B. J. (2018). Ventral striatal response during decision making involving risk and reward is associated with future binge drinking in adolescents. Neuropsychopharmacology, 43(9), 1884–1890. doi: 10.1038/s41386-018-0087-8

39. Must, A., & Anderson, S. E. (2006). Body mass index in children and adolescents: considerations for population-based applications. Int J Obes (Lond), 30(4), 590–594. doi: 10.1038/sj.ijo.0803300

40. Pauli, W. M., Nili, A. N., & Tyszka, J. M. (2018). A high-resolution probabilistic in vivo atlas of human subcortical brain nuclei. Sci Data, 5, 180063. doi: 10.1038/sdata.2018.63

41. Perlaki, G., Molnar, D., Smeets, P. A. M., Ahrens, W., Wolters, M., Eiben, G., … Orsi, G. (2018). Volumetric gray matter measures of amygdala and accumbens in childhood overweight/obesity. PLoS One, 13(10), e0205331. doi: 10.1371/journal.pone.0205331

42. Petersen, A. C., Crockett, L., Richards, M., & Boxer, A. (1988). A self-report measure of pubertal status: Reliability, validity, and initial norms. J Youth Adolesc, 17(2), 117–133. doi: 10.1007/bf01537962

43. Phillips, R. G., & LeDoux, J. E. (1992). Differential contribution of amygdala and hippocampus to cued and contextual fear conditioning. Behav Neurosci, 106(2), 274–285.

44. Pitkanen, A., & Amaral, D. G. (1998). Organization of the intrinsic connections of the monkey amygdaloid complex: projections originating in the lateral nucleus. J Comp Neurol, 398(3), 431-458. Retrieved from https://onlinelibrary.wiley.com/doi/pdf/10.1002/%28SICI%291096-9861%2819980831%29398%3A3%3C431%3A%3AAID-CNE9%3E3.0.CO%3B2-0

45. Raznahan, A., Lerch, J. P., Lee, N., Greenstein, D., Wallace, G. L., Stockman, M., … Giedd, J. N. (2011). Patterns of coordinated anatomical change in human cortical development: a longitudinal neuroimaging study of maturational coupling. Neuron, 72(5), 873–884. doi: 10.1016/j.neuron.2011.09.028

46. Rollins, B. L., & King, B. M. (2000). Amygdala-lesion obesity: what is the role of the various amygdaloid nuclei? Am J Physiol Regul Integr Comp Physiol, 279(4), R1348–1356. doi: 10.1152/ajpregu.2000.279.4.R1348

47. Sah, P., Faber, E. S., Lopez De Armentia, M., & Power, J. (2003). The amygdaloid complex: anatomy and physiology. Physiol Rev, 83(3), 803–834. doi: 10.1152/physrev.00002.2003

48. Sananes, C. B., & Davis, M. (1992). N-methyl-D-aspartate lesions of the lateral and basolateral nuclei of the amygdala block fear-potentiated startle and shock sensitization of startle. Behav Neurosci, 106(1), 72–80.

49. Saygin, Z. M., Kliemann, D., Iglesias, J. E., van der Kouwe, A. J. W., Boyd, E., Reuter, M., … Augustinack, J. C. (2017). High-resolution magnetic resonance imaging reveals nuclei of the human amygdala: manual segmentation to automatic atlas. Neuroimage, 155, 370–382. doi: 10.1016/j.neuroimage.2017.04.046

50. Scheuer, H., Alarcon, G., Demeter, D. V., Earl, E., Fair, D. A., & Nagel, B. J. (2017). Reduced fronto-amygdalar connectivity in adolescence is associated with increased depression symptoms over time. Psychiatry Res Neuroimaging, 266, 35–41. doi: 10.1016/j.pscychresns.2017.05.012

51. Schoenbaum, G., Chiba, A. A., & Gallagher, M. (1999). Neural encoding in orbitofrontal cortex and basolateral amygdala during olfactory discrimination learning. J Neurosci, 19(5), 1876-1884. Retrieved from http://www.jneurosci.org/content/jneuro/19/5/1876.full.pdf

52. Shaffer, D., Fisher, P., Lucas, C. P., Dulcan, M. K., & Schwab-Stone, M. E. (2000). NIMH Diagnostic Interview Schedule for Children Version IV (NIMH DISC-IV): description, differences from previous versions, and reliability of some common diagnoses. J Am Acad Child Adolesc Psychiatry, 39(1), 28–38. doi: 10.1097/00004583-200001000-00014

53. Smith, S. M., Jenkinson, M., Woolrich, M. W., Beckmann, C. F., Behrens, T. E., Johansen-Berg, H., … Matthews, P. M. (2004). Advances in functional and structural MR image analysis and implementation as FSL. Neuroimage, 23 Suppl 1, S208–219. doi: 10.1016/j.neuroimage.2004.07.051

54. Solano-Castiella, E., Schafer, A., Reimer, E., Turke, E., Proger, T., Lohmann, G., … Turner, R. (2011). Parcellation of human amygdala in vivo using ultra high field structural MRI. Neuroimage, 58(3), 741–748. doi: 10.1016/j.neuroimage.2011.06.047

55. Stan, A. D., Ghose, S., Gao, X. M., Roberts, R. C., Lewis-Amezcua, K., Hatanpaa, K. J., & Tamminga, C. A. (2006). Human postmortem tissue: what quality markers matter? Brain Res, 1123(1), 1–11. doi: 10.1016/j.brainres.2006.09.025

56. Tosevski, J., Malikovic, A., Mojsilovic-Petrovic, J., Lackovic, V., Peulic, M., Sazdanovic, P., & Alexopulos, C. (2002). Types of neurons and some dendritic patterns of basolateral amygdala in humans--a golgi study. Ann Anat, 184(1), 93–103. doi: 10.1016/s0940-9602(02)80042-5

57. Tottenham, N., & Gabard-Durnam, L. J. (2017). The developing amygdala: a student of the world and a teacher of the cortex. Curr Opin Psychol, 17, 55–60. doi: 10.1016/j.copsyc.2017.06.012

58. Tustison, N. J., Avants, B. B., Cook, P. A., Zheng, Y., Egan, A., Yushkevich, P. A., & Gee, J. C. (2010). N4ITK: improved N3 bias correction. IEEE Trans Med Imaging, 29(6), 1310–1320. doi: 10.1109/tmi.2010.2046908

59. Tyszka, J. M., & Pauli, W. M. (2016). In vivo delineation of subdivisions of the human amygdaloid complex in a high-resolution group template. Hum Brain Mapp, 37(11), 3979–3998. doi: 10.1002/hbm.23289

60. Wan, F. J., & Swerdlow, N. R. (1997). The basolateral amygdala regulates sensorimotor gating of acoustic startle in the rat. Neuroscience, 76(3), 715–724.

61. Wierenga, L., Langen, M., Ambrosino, S., van Dijk, S., Oranje, B., & Durston, S. (2014). Typical development of basal ganglia, hippocampus, amygdala and cerebellum from age 7 to 24. Neuroimage, 96, 67–72. doi: 10.1016/j.neuroimage.2014.03.072

62. Wierenga, L. M., Bos, M. G. N., Schreuders, E., Vd Kamp, F., Peper, J. S., Tamnes, C. K., & Crone, E. A. (2018). Unraveling age, puberty and testosterone effects on subcortical brain development across adolescence. Psychoneuroendocrinology, 91, 105–114. doi: 10.1016/j.psyneuen.2018.02.034

63. Woolrich, M. W., Jbabdi, S., Patenaude, B., Chappell, M., Makni, S., Behrens, T., … Smith, S. M. (2009). Bayesian analysis of neuroimaging data in FSL. Neuroimage, 45(1 Suppl), S173–186. doi: 10.1016/j.neuroimage.2008.10.055

