## Supplemental Information for "Restructuring of amygdala subregion apportion across adolescence"

### Supporting Information

#### Analysis of Relative Volume Fraction (RVF) covarying for Intracranial Volume (ICV)

Statistical Analysis

The following are the Generalized Additive Mixed Models (GAMMs) for the RVF plus ICV to assess age, sex, and age-by-sex interaction while controlling for Hemisphere, BMIz, ICV, and SES:

$$Supporting\ Model\ (SM)\ 1: RVF + ICV_{ij} = \beta_0 + s_1(Age_i) + \beta_1 Male_i + s_2(Age_i) \times Male_i + \beta_2 Hemisphere_{ij} + \beta_3 BMIz_i + \beta_4 ICV_i + \beta_5 SES_i + U_i + \varepsilon_{ij}$$

Next, the following are the GAMMs for the RVF plus ICV to assess whether PDS is a better predictor of volumetric associations than age while controlling for Hemisphere, BMIz, ICV, and SES:

$$SM2: RVF + ICV_{ij} = \beta_0 + s_1(Age_i) + \beta_1 Hemisphere_{ij} + \beta_2 BMIz_i + \beta_3 ICV_i + U_i + \varepsilon_{ij}$$

$$SM3: RVF + ICV_{ij} = \beta_0 + s_1(Pubertal\ Stage_i) + \beta_1 Hemisphere_{ij} + \beta_2 BMIz_i + \beta_3 ICV_i + U_i + \varepsilon_{ij}$$

$$SM4: RVF + ICV_{ij} = \beta_0 + s_1(Age_i) + s_2(Pubertal\ Stage_i) + s_3(Age_i, Pubertal\ Stage_i) + \beta_2 Hemisphere_{ij} + \beta_3 BMIz_i + \beta_4 ICV_i + U_i + \varepsilon_{ij}$$

Akaike Information Criterion (AIC) and Likelihood ratio tests ($p < 0.05$) were used to compare model fits. To reduce type I error, each set of models across the 9 ROIs were corrected for multiple comparisons using the Bonferroni correction method (Bonferroni, 1936), with p-values <0.0056 deemed significant.

Results

*Age and sex differences in amygdala composition:*

GAMM results (SM1) examining the associations between amygdala subregion RVF with age, sex, hemisphere, BMIz, ICV, SES, and age-by-sex interactions are presented in **Supporting Table 2**. A significant age-by-sex interaction was detected for the LA (Adj $R^2$=.06), BLVPL (Adj $R^2$=.13), CEN (Adj $R^2$=.09), ATA (Adj $R^2$=.12), and ASTA (Adj $R^2$=.06), which are equivalent to the results presented for RVF without ICV added (**Table 5**). The relative volumes of the BLDI, BM, CMN, and AAA did not relate to age, sex, or their interaction.

*Pubertal development and amygdala composition in males and females:*

GAMM model outputs for age (SM2), puberty (SM3), and age-by-puberty (SM4) for each RVF in each sex separately are presented in **Supporting Table 5** for males and **Supporting Table 6** for females. For females, no significant age, puberty, or age-by-puberty associations were seen for any of the 9 amygdala subregions. In males, age was again found to be significantly associated with RVFs of the BLVPL, CEN, ATA, and ASTA (SM2: p's $\leq$ 0.0056), and trending for LA (SM2: p=0.0068). In addition, puberty was found to significantly relate to RVFs of the BLVPL, CEN, and ATA (SM3: p's $\leq$ 0.0056). There were no age-by-pubertal interactions that were significant for any of the 9 amygdala subregions. For the BLVPL and the CEN, best-fit model comparisons showed that the age and puberty model was significantly better than the model including only puberty (SM3 vs. SM4: p's<0.05), but was no significantly better than age alone (SM2 vs. SM4: p's>0.05); this indicates that age is a better predictor for BLVPL and CEN RVFs – while controlling for ICV – than puberty. The ASTA was trending for the age model to be the best fit (p=0.0875) and showed a smaller AIC and BIC with a larger log-likelihood indicating a better fit model compared to both the PDS only model and the interaction model. In contract, for the ATA, both age alone and puberty alone are significantly different from the age-by-puberty model, with AIC and log-likelihood indicating puberty was a better fit for the ATA in males compared to age.

**Supporting Information**

### Tables

**Supporting Table 1.** *Amygdala Subregion ROIs.*

| | CIT168 Regions | | | | | | | | |
|---|---|---|---|---|---|---|---|---|---|
| *Subregion name* | Lateral nucleus | Basolateral dorsal and intermediate subdivision | Basolateral ventral and paralaminar subdivision | Basomedial nucleus | Cortical and medial nuclei | Central nucleus | Anterior amygdala area | Amygdala transition area | Amygdalostriatal transition area |
| *Subregion abbreviation* | (LA) | (BLDI) | (BLVPL) | (BM) | (CMN) | (CEN) | (AAA) | (ATA) | (ASTA) |
| *Subregion location or other common names* | Surrounded ventrally and caudally by lateral ventricle, and laterally by temporal lobe white matter; primary input from neocortex | Subdivision of the basolateral nucleus; the basolateral nucleus lies medially to the lateral nucleus (LA) | Subdivision of the basolateral nucleus; the basolateral nucleus lies medially to the lateral nucleus (LA) | Ventrally bounded by BLV; BM also known as the accessory basal nucleus | Lies along dorsomedial surface of amygdaloid complex | Major output nuclei; lies dorsally and caudally within complex | Lies rostrally and caudally within complex; borders pariamygdaloid claustrum and basolateral complex | Boundary between enterhinal cortex and CMN | Lies medially and ventrally to temporal branch of anterior commissure; borders ventral putamen |
| *Subregion content* | LA exclusively | Merger between the basolateral nucleus' 2 out of 3 divisions: dorsal (BLD) and intermediate (BLI) | Merger between the basolateral nucleus' 3rd division, ventral (BLV), and the paralaminar nucleus | BM exclusively | Merger between cortical nucleus (CoA) and corticomedical group (CoMe); CoMe is made up of the internal boundaries between CoA, posterior cortical nucleus (CoP), amygdalohippocampal (AHA), nucleus of the lateral olfactory tract (NLOT), and medial nucleus (Me) | CEN exclusively | AAA exclusively | Merger between amygdalocortical and amygdalohippocampal transition areas, and periamygdaloid cortex | ASTA exclusively |

Descriptions based on the CIT168 atlas by Tyszka JM, Pauli WM (2016)

**Supporting Table 2.** *GAMM results for amygdala subregion Relative Volume Fraction (RVF) associations with age, sex, and age sex interaction, controlling for hemisphere, BMI, and ICV.*

| **LA** | | | | | |
|---|---|---|---|---|---|
| | *edf* | *Ref.df* | *F* | *p-value* | Adj R squared |
| s(age) | 9.39E-03 | 3 | 0 | 0.85 | |
| s(age*sex(male)) | 1.248401 | 3 | 2.68 | ***0.00346*** | |
| | *Estimate* | *SE* | *t-value* | *p-value* | |
| Intercept | 0.2140 | 0.0063 | 33.81 | ***7.23E-157*** | |
| Sex (male) | 0.0009 | 0.0012 | 0.75 | 0.45 | 0.0587 |
| Hemisphere (right) | 0.0052 | 0.0006 | 8.82 | ***6.68E-18*** | |
| BMI | 0.0006 | 0.0005 | 1.11 | 0.27 | |
| ICV | -2.31E-09 | 4.49E-09 | -0.51 | 0.61 | |
| SES | -1.42E-05 | 3.69E-05 | -0.38 | 0.70 | |

| **CEN** | | | | | |
|---|---|---|---|---|---|
| | *edf* | *Ref.df* | *F* | *p-value* | Adj R squared |
| s(age) | 0.65 | 3 | 0.45 | 0.13 | |
| s(age*sex(male)) | 2.17 | 3 | 3.69 | ***0.00105*** | |
| | *Estimate* | *SE* | *t-value* | *p-value* | |
| Intercept | 0.0298 | 0.0011 | 26.66 | ***8.49E-113*** | |
| Sex (male) | -0.0007 | 0.0002 | -3.26 | ***1.14E-03*** | 0.0927 |
| Hemisphere (right) | -0.0008 | 0.0001 | -7.07 | ***3.38E-12*** | |
| BMI | 0.0001 | 0.0001 | 1.42 | 0.16 | |
| ICV | 3.41E-10 | 7.93E-10 | 0.43 | 0.67 | |
| SES | 4.45E-06 | 6.52E-06 | 0.68 | 0.50 | |

| **BLDI** | | | | | |
|---|---|---|---|---|---|
| | *edf* | *Ref.df* | *F* | *p-value* | Adj R squared |
| s(age) | 0 | 3 | 0 | 0.33 | |
| s(age*sex(male)) | 0 | 3 | 0 | 0.21 | |
| | *Estimate* | *SE* | *t-value* | *p-value* | |
| Intercept | 0.1266 | 0.0020 | 64.33 | ***1.74E-320*** | |
| Sex (male) | 3.64E-06 | 0.0004 | 0.01 | 0.99 | 0.1471 |
| Hemisphere (right) | -0.0030 | 0.0002 | -16.59 | ***2.08E-53*** | |
| BMI | -0.0002 | 0.0002 | -1.06 | 0.29 | |
| ICV | 1.18E-09 | 1.40E-09 | 0.84 | 0.40 | |
| SES | -1.55E-05 | 1.15E-05 | -1.35 | 0.18 | |

| **AAA** | | | | | |
|---|---|---|---|---|---|
| | *edf* | *Ref.df* | *F* | *p-value* | Adj R squared |
| s(age) | 0.12 | 3 | 0.05 | 0.29 | |
| s(age*sex(male)) | 0 | 3 | 0 | 0.64 | |
| | *Estimate* | *SE* | *t-value* | *p-value* | |
| Intercept | 0.0417 | 0.0013 | 31.83 | ***8.01E-145*** | |
| Sex (male) | -4.80E-05 | 0.0003 | -0.19 | 0.85 | 0.0058 |
| Hemisphere (right) | -0.0003 | 0.0002 | -1.91 | 0.06 | |
| BMI | 0.0001 | 0.0001 | 0.57 | 0.57 | |
| ICV | -1.52E-09 | 9.31E-10 | -1.63 | 0.10 | |
| SES | 3.33E-06 | 7.64E-06 | 0.44 | 0.66 | |

| **BLVPL** | | | | | |
|---|---|---|---|---|---|
| | *edf* | *Ref.df* | *F* | *p-value* | Adj R squared |
| s(age) | 0 | 3 | 0 | 0.71 | |
| s(age*sex(male)) | 2.38 | 3 | 12.06 | ***7.46E-09*** | |
| | *Estimate* | *SE* | *t-value* | *p-value* | |
| Intercept | 0.0756 | 0.0019 | 39.43 | ***1.99E-190*** | |
| Sex (male) | 0.0008 | 0.0004 | 2.17 | 0.03 | 0.1318 |
| Hemisphere (right) | -0.0026 | 0.0003 | -8.53 | ***7.24E-17*** | |
| BMI | -0.0003 | 0.0002 | -1.68 | 0.09 | |
| ICV | 1.76E-09 | 1.36E-09 | 1.30 | 0.19 | |
| SES | -1.88E-05 | 1.12E-05 | -1.69 | 0.09 | |

| **ATA** | | | | | |
|---|---|---|---|---|---|
| | *edf* | *Ref.df* | *F* | *p-value* | Adj R squared |
| s(age) | 0 | 3 | 0 | 0.56 | |
| s(age*sex(male)) | 2.26 | 3 | 15.74 | ***1.38E-11*** | |
| | *Estimate* | *SE* | *t-value* | *p-value* | |
| Intercept | 0.0603 | 0.0023 | 26.75 | ***2.14E-113*** | |
| Sex (male) | 0.0014 | 0.0004 | 3.19 | ***1.49E-03*** | 0.1197 |
| Hemisphere (right) | 0.0014 | 0.0002 | 6.51 | ***1.29E-10*** | |
| BMI | -0.0003 | 0.0002 | -1.75 | 0.08 | |
| ICV | -5.60E-10 | 1.60E-09 | -0.35 | 0.73 | |
| SES | -1.01E-05 | 1.32E-05 | -0.77 | 0.44 | |

| **BM** | | | | | |
|---|---|---|---|---|---|
| | *edf* | *Ref.df* | *F* | *p-value* | Adj R squared |
| s(age) | 0 | 3 | 0 | 0.67 | |
| s(age*sex(male)) | 0 | 3 | 0 | 0.38 | |
| | *Estimate* | *SE* | *t-value* | p-value | |
| Intercept | 0.0681 | 0.0014 | 48.02 | ***3.60E-239*** | |
| Sex (male) | -0.0007 | 0.0003 | -2.53 | 0.01 | 0.0572 |
| Hemisphere (right) | -0.0014 | 0.0002 | -6.16 | ***1.13E-09*** | |
| BMI | -0.0002 | 0.0001 | -1.46 | 0.14 | |
| ICV | 3.63E-09 | 1.01E-09 | 3.61 | ***3.26E-04*** | |
| SES | 3.19E-06 | 8.25E-06 | 0.39 | 0.70 | |

| **ASTA** | | | | | |
|---|---|---|---|---|---|
| | *edf* | *Ref.df* | *F* | *p-value* | Adj R squared |
| s(age) | 0 | 3 | 0 | 0.31 | |
| s(age*sex(male)) | 1.34 | 3 | 2.55 | ***0.00556*** | |
| | *Estimate* | *SE* | *t-value* | *p-value* | |
| Intercept | 0.0440 | 0.0016 | 26.98 | ***7.57E-115*** | |
| Sex (male) | -0.0010 | 0.0003 | -3.20 | ***1.45E-03*** | 0.0553 |
| Hemisphere (right) | -0.0005 | 0.0002 | -3.40 | ***7.04E-04*** | |
| BMI | 0.0002 | 0.0001 | 1.14 | 0.25 | |
| ICV | -3.96E-10 | 1.16E-09 | -0.34 | 0.73 | |
| SES | 2.08E-05 | 9.51E-06 | 2.19 | 0.03 | |

| **CMN** | | | | | |
|---|---|---|---|---|---|
| | *edf* | *Ref.df* | *F* | *p-value* | Adj R squared |
| s(age) | 0 | 3 | 0 | 0.56 | |
| s(age*sex(male)) | 0.08 | 3 | 0.03 | 0.28 | |
| | *Estimate* | *SE* | *t-value* | *p-value* | |
| Intercept | 0.1105 | 0.0029 | 38.08 | ***1.09E-182*** | |
| Sex (male) | -0.0002 | 0.0006 | -0.39 | 0.70 | 0.0174 |
| Hemisphere (right) | -0.0015 | 0.0003 | -5.58 | ***3.23E-08*** | |
| BMI | 8.10E-05 | 0.0002 | 0.34 | 0.74 | |
| ICV | -1.44E-09 | 2.06E-09 | -0.70 | 0.49 | |
| SES | 1.17E-05 | 1.69E-05 | 0.69 | 0.49 | |

Notes: In each model, for the parametric terms, the estimate, standard error (SE), t-value, and p-value are shown, for the smooth terms, the estimated degree of freedom (edf), reference degree of freedom (Ref.df), F-score, and p-value are shown; the Adjusted $R^2$ for each model is also shown. P-values of significance level less than 0.0056 bolded. Abbreviations: See Table 1.

**Supporting Table 3.** *GAMM amygdala subregion Absolute Probabilistic Estimate results for age, pubertal status, and age-by-pubertal status interaction for females*

| *FEMALES* | | | | | | | | | | | | | |
|---|---|---|---|---|---|---|---|---|---|---|---|---|---|
| **Total** | **Smooth Terms** | | | | | | **Model Fit** | | | | | | |
| | **Terms** | **edf** | **Ref.df** | **F** | **p-value** | **R2** | **df** | **AIC** | **BIC** | **logLik** | **Test** | **L.Ratio** | **p-value** |
| M4 | s(age) | 0.03 | 3.00 | 0.00 | 0.5761 | 0.3851 | 8 | 4152.45 | 4183.36 | -2068.22 | M4 vs. M6 | 0.09 | 0.999 |
| M5 | s(pds) | 0.04 | 3.00 | 0.00 | 0.7369 | 0.3850 | 8 | 4152.47 | 4183.37 | -2068.23 | M5 vs. M6 | 0.11 | 0.999 |
| M6 | ti(age) | 0.01 | 3.00 | 0.00 | 0.5024 | 0.3836 | 12 | 4160.36 | 4206.72 | -2068.18 | | | |
| | ti(pds) | 0.00 | 3.00 | 0.00 | 0.8064 | | | | | | | | |
| | ti(age, pds) | 1.00 | 1.00 | 0.08 | 0.7829 | | | | | | | | |
| **LA** | **Smooth Terms** | | | | | | **Model Fit** | | | | | | |
| | **Terms** | **edf** | **Ref.df** | **F** | **p-value** | **R2** | **df** | **AIC** | **BIC** | **logLik** | **Test** | **L.Ratio** | **p-value** |
| M4 | s(age) | 0.15 | 3.00 | 0.04 | 0.3584 | 0.2957 | 8 | 3316.79 | 3347.70 | -1650.39 | M4 vs. M6 | 0.28 | 0.991 |
| M5 | s(pds) | 0.03 | 3.00 | 0.00 | 0.8242 | 0.2952 | 8 | 3316.79 | 3347.70 | -1650.39 | M5 vs. M6 | 0.28 | 0.991 |
| M6 | ti(age) | 0.01 | 3.00 | 0.00 | 0.3939 | 0.2940 | 12 | 3324.51 | 3370.87 | -1650.25 | | | |
| | ti(pds) | 0.00 | 3.00 | 0.00 | 1.0000 | | | | | | | | |
| | ti(age, pds) | 1.00 | 1.00 | 0.25 | 0.6184 | | | | | | | | |
| **BLDI** | **Smooth Terms** | | | | | | **Model Fit** | | | | | | |
| | **Terms** | **edf** | **Ref.df** | **F** | **p-value** | **R2** | **df** | **AIC** | **BIC** | **logLik** | **Test** | **L.Ratio** | **p-value** |
| M4 | s(age) | 0.01 | 3.00 | 0.00 | 0.6118 | 0.3318 | 8 | 2856.33 | 2887.24 | -1420.16 | M4 vs. M6 | 0.01 | 1.000 |
| M5 | s(pds) | 0.00 | 3.00 | 0.00 | 0.8412 | 0.3318 | 8 | 2856.33 | 2887.24 | -1420.16 | M5 vs. M6 | 0.01 | 1.000 |
| M6 | ti(age) | 0.01 | 3.00 | 0.00 | 0.5574 | 0.3299 | 12 | 2864.32 | 2910.68 | -1420.16 | | | |
| | ti(pds) | 0.00 | 3.00 | 0.00 | 0.8359 | | | | | | | | |
| | ti(age, pds) | 1.00 | 1.00 | 0.01 | 0.9278 | | | | | | | | |
| **BLVPL** | **Smooth Terms** | | | | | | **Model Fit** | | | | | | |
| | **Terms** | **edf** | **Ref.df** | **F** | **p-value** | **R2** | **df** | **AIC** | **BIC** | **logLik** | **Test** | **L.Ratio** | **p-value** |
| M4 | s(age) | 0.00 | 3.00 | 0.00 | 0.6116 | 0.2702 | 8 | 2615.46 | 2646.37 | -1299.73 | M4 vs. M6 | 0.51 | 0.973 |
| M5 | s(pds) | 0.06 | 3.00 | 0.01 | 0.4929 | 0.2703 | 8 | 2615.50 | 2646.41 | -1299.75 | M5 vs. M6 | 0.54 | 0.969 |
| M6 | ti(age) | 0.00 | 3.00 | 0.00 | 0.5115 | 0.2699 | 12 | 2622.96 | 2669.32 | -1299.48 | | | |
| | ti(pds) | 0.00 | 3.00 | 0.00 | 0.7007 | | | | | | | | |
| | ti(age, pds) | 1.00 | 1.00 | 0.51 | 0.4786 | | | | | | | | |
| **BM** | **Smooth Terms** | | | | | | **Model Fit** | | | | | | |
| | **Terms** | **edf** | **Ref.df** | **F** | **p-value** | **R2** | **df** | **AIC** | **BIC** | **logLik** | **Test** | **L.Ratio** | **p-value** |
| M4 | s(age) | 0.01 | 3.00 | 0.00 | 0.7155 | 0.3290 | 8 | 2472.62 | 2503.53 | -1228.31 | M4 vs. M6 | 0.95 | 0.917 |
| M5 | s(pds) | 0.01 | 3.00 | 0.00 | 0.5035 | 0.3291 | 8 | 2472.62 | 2503.53 | -1228.31 | M5 vs. M6 | 0.95 | 0.918 |
| M6 | ti(age) | 0.00 | 3.00 | 0.00 | 0.7619 | 0.3347 | 12 | 2479.67 | 2526.03 | -1227.84 | | | |
| | ti(pds) | 0.01 | 3.00 | 0.00 | 0.4928 | | | | | | | | |
| | ti(age, pds) | 1.62 | 1.62 | 1.29 | 0.4065 | | | | | | | | |
| **CMN** | **Smooth Terms** | | | | | | **Model Fit** | | | | | | |
| | **Terms** | **edf** | **Ref.df** | **F** | **p-value** | **R2** | **df** | **AIC** | **BIC** | **logLik** | **Test** | **L.Ratio** | **p-value** |
| M4 | s(age) | 0.01 | 3.00 | 0.00 | 0.7224 | 0.2627 | 8 | 2739.20 | 2770.10 | -1361.60 | M4 vs. M6 | 0.69 | 0.953 |
| M5 | s(pds) | 0.03 | 3.00 | 0.00 | 0.7139 | 0.2626 | 8 | 2739.22 | 2770.13 | -1361.61 | M5 vs. M6 | 0.71 | 0.950 |
| M6 | ti(age) | 0.01 | 3.00 | 0.00 | 0.6127 | 0.2629 | 12 | 2746.51 | 2792.87 | -1361.25 | | | |
| | ti(pds) | 0.00 | 3.00 | 0.00 | 1.0000 | | | | | | | | |
| | ti(age, pds) | 1.00 | 1.00 | 0.68 | 0.4109 | | | | | | | | |
| **CEN** | **Smooth Terms** | | | | | | **Model Fit** | | | | | | |
| | **Terms** | **edf** | **Ref.df** | **F** | **p-value** | **R2** | **df** | **AIC** | **BIC** | **logLik** | **Test** | **L.Ratio** | **p-value** |
| M4 | s(age) | 0.02 | 3.00 | 0.00 | 0.4642 | 0.2846 | 8 | 1935.08 | 1965.99 | -959.54 | M4 vs. M6 | 0.66 | 0.956 |
| M5 | s(pds) | 0.16 | 3.00 | 0.06 | 0.2781 | 0.2854 | 8 | 1935.06 | 1965.97 | -959.53 | M5 vs. M6 | 0.64 | 0.959 |
| M6 | ti(age) | 0.00 | 3.00 | 0.00 | 0.5891 | 0.2846 | 12 | 1942.42 | 1988.79 | -959.21 | | | |
| | ti(pds) | 0.00 | 3.00 | 0.00 | 0.4246 | | | | | | | | |
| | ti(age, pds) | 1.00 | 1.00 | 0.65 | 0.4236 | | | | | | | | |
| **AAA** | **Smooth Terms** | | | | | | **Model Fit** | | | | | | |
| | **Terms** | **edf** | **Ref.df** | **F** | **p-value** | **R2** | **df** | **AIC** | **BIC** | **logLik** | **Test** | **L.Ratio** | **p-value** |
| M4 | s(age) | 0.09 | 3.00 | 0.02 | 0.4033 | 0.1937 | 8 | 2232.85 | 2263.76 | -1108.42 | M4 vs. M6 | 0.60 | 0.963 |
| M5 | s(pds) | 0.02 | 3.00 | 0.00 | 0.5583 | 0.1934 | 8 | 2232.84 | 2263.75 | -1108.42 | M5 vs. M6 | 0.59 | 0.964 |
| M6 | ti(age) | 0.01 | 3.00 | 0.00 | 0.4909 | 0.1931 | 12 | 2240.24 | 2286.61 | -1108.12 | | | |
| | ti(pds) | 0.00 | 3.00 | 0.00 | 0.8013 | | | | | | | | |
| | ti(age, pds) | 1.00 | 1.00 | 0.57 | 0.4502 | | | | | | | | |
| **ATA** | **Smooth Terms** | | | | | | **Model Fit** | | | | | | |
| | **Terms** | **edf** | **Ref.df** | **F** | **p-value** | **R2** | **df** | **AIC** | **BIC** | **logLik** | **Test** | **L.Ratio** | **p-value** |
| M4 | s(age) | 1.13 | 3.00 | 0.50 | 0.2561 | 0.2479 | 8 | 2445.68 | 2476.59 | -1214.84 | M4 vs. M6 | 4.10 | 0.393 |
| M5 | s(pds) | 0.43 | 3.00 | 0.24 | 0.1962 | 0.2445 | 8 | 2444.79 | 2475.69 | -1214.39 | M5 vs. M6 | 3.21 | 0.524 |
| M6 | ti(age) | 0.01 | 3.00 | 0.00 | 0.5610 | 0.2506 | 12 | 2449.58 | 2495.94 | -1212.79 | | | |
| | ti(pds) | 0.00 | 3.00 | 0.00 | 0.6691 | | | | | | | | |
| | ti(age, pds) | 1.00 | 1.00 | 3.34 | 0.0681 | | | | | | | | |
| **ASTA** | **Smooth Terms** | | | | | | **Model Fit** | | | | | | |
| | **Terms** | **edf** | **Ref.df** | **F** | **p-value** | **R2** | **df** | **AIC** | **BIC** | **logLik** | **Test** | **L.Ratio** | **p-value** |
| M4 | s(age) | 0.01 | 3.00 | 0.00 | 0.7962 | 0.2753 | 8 | 2231.40 | 2262.31 | -1107.70 | M4 vs. M6 | 0.21 | 0.995 |
| M5 | s(pds) | 0.06 | 3.00 | 0.01 | 0.5088 | 0.2754 | 8 | 2231.43 | 2262.33 | -1107.71 | M5 vs. M6 | 0.23 | 0.994 |
| M6 | ti(age) | 0.00 | 3.00 | 0.00 | 0.8307 | 0.2739 | 12 | 2239.19 | 2285.56 | -1107.60 | | | |
| | ti(pds) | 0.00 | 3.00 | 0.00 | 0.6044 | | | | | | | | |
| | ti(age, pds) | 1.00 | 1.00 | 0.20 | 0.6590 | | | | | | | | |

Notes: In each model, the smooth terms, the estimated degree of freedom (edf), reference degree of freedom (Ref.df), F-score, p-value, and Adjusted $R^2$ for each model is shown; p-value < 0.0056 bolded (Bonferroni corrected). Between model comparisons include the df, AIC, log-likelihood ratio (L Ratio) and p-values < 0.05 bolded.

**Supporting Table 4.** *GAMM amygdala subregion Relative Volume Fraction (RVF) results for age, pubertal status, and age-by-pubertal status interaction for females*

| FEMALES | | | | | | | | | | | | | |
|---|---|---|---|---|---|---|---|---|---|---|---|---|---|
| **LA** | **Smooth Terms** | | | | | | **Model Fit** | | | | | | |
| | **Terms** | **edf** | **Ref.df** | **F** | **p-value** | **R2** | **df** | **AIC** | **BIC** | **logLik** | **Test** | **L.Ratio** | **p-value** |
| M7 | s(age) | 0.21 | 3.00 | 0.09 | 0.2727 | 0.0380 | 7 | -2169.36 | -2142.31 | 1091.68 | M7 vs. M9 | 2.21 | 0.697 |
| M8 | s(pds) | 0.25 | 3.00 | 0.11 | 0.2495 | 0.0382 | 7 | -2169.37 | -2142.33 | 1091.69 | M8 vs. M9 | 2.19 | 0.700 |
| | ti(age) | 0.00 | 3.00 | 0.00 | 0.5066 | | | | | | | | |
| M9 | ti(pds) | 0.00 | 3.00 | 0.00 | 0.6507 | 0.0406 | 11 | -2163.57 | -2121.07 | 1092.78 | | | |
| | ti(age, pds) | 1.00 | 1.00 | 2.21 | 0.1376 | | | | | | | | |
| **BLDI** | **Smooth Terms** | | | | | | **Model Fit** | | | | | | |
| | **Terms** | **edf** | **Ref.df** | **F** | **p-value** | **R2** | **df** | **AIC** | **BIC** | **logLik** | **Test** | **L.Ratio** | **p-value** |
| M7 | s(age) | 0.02 | 3.00 | 0.00 | 1.0000 | 0.1659 | 7 | -3016.31 | -2989.26 | 1515.15 | M7 vs. M9 | 0.16 | 0.997 |
| M8 | s(pds) | 0.02 | 3.00 | 0.00 | 1.0000 | 0.1659 | 7 | -3016.31 | -2989.26 | 1515.15 | M8 vs. M9 | 0.16 | 0.997 |
| | ti(age) | 0.00 | 3.00 | 0.00 | 1.0000 | | | | | | | | |
| M9 | ti(pds) | 0.00 | 3.00 | 0.00 | 0.8622 | 0.1641 | 11 | -3008.47 | -2965.97 | 1515.24 | | | |
| | ti(age, pds) | 1.00 | 1.00 | 0.15 | 0.6982 | | | | | | | | |
| **BLVPL** | **Smooth Terms** | | | | | | **Model Fit** | | | | | | |
| | **Terms** | **edf** | **Ref.df** | **F** | **p-value** | **R2** | **df** | **AIC** | **BIC** | **logLik** | **Test** | **L.Ratio** | **p-value** |
| M7 | s(age) | 0.01 | 3.00 | 0.00 | 0.6673 | 0.0896 | 7 | -2874.69 | -2847.65 | 1444.35 | M7 vs. M9 | 3.34 | 0.503 |
| M8 | s(pds) | 0.03 | 3.00 | 0.01 | 0.3174 | 0.0897 | 7 | -2874.70 | -2847.65 | 1444.35 | M8 vs. M9 | 3.33 | 0.504 |
| | ti(age) | 1.84 | 3.00 | 2.20 | 0.0156 | | | | | | | | |
| M9 | ti(pds) | 0.00 | 3.00 | 0.00 | 0.8456 | 0.1302 | 11 | -2870.03 | -2827.53 | 1446.02 | | | |
| | ti(age, pds) | 3.86 | 3.86 | 2.72 | 0.0455 | | | | | | | | |
| **BM** | **Smooth Terms** | | | | | | **Model Fit** | | | | | | |
| | **Terms** | **edf** | **Ref.df** | **F** | **p-value** | **R2** | **df** | **AIC** | **BIC** | **logLik** | **Test** | **L.Ratio** | **p-value** |
| M7 | s(age) | 0.00 | 3.00 | 0.00 | 0.4684 | 0.0443 | 7 | -3126.05 | -3099.00 | 1570.02 | M7 vs. M9 | 1.60 | 0.809 |
| M8 | s(pds) | 0.02 | 3.00 | 0.01 | 0.2960 | 0.0444 | 7 | -3126.05 | -3099.00 | 1570.02 | M8 vs. M9 | 1.60 | 0.810 |
| | ti(age) | 0.00 | 3.00 | 0.00 | 0.7049 | | | | | | | | |
| M9 | ti(pds) | 0.00 | 3.00 | 0.00 | 0.6353 | 0.0459 | 11 | -3119.64 | -3077.14 | 1570.82 | | | |
| | ti(age, pds) | 1.00 | 1.00 | 1.59 | 0.2096 | | | | | | | | |
| **CMN** | **Smooth Terms** | | | | | | **Model Fit** | | | | | | |
| | **Terms** | **edf** | **Ref.df** | **F** | **p-value** | **R2** | **df** | **AIC** | **BIC** | **logLik** | **Test** | **L.Ratio** | **p-value** |
| M7 | s(age) | 0.00 | 3.00 | 0.00 | 0.6994 | 0.0146 | 7 | -2724.66 | -2697.61 | 1369.33 | M7 vs. M9 | 1.46 | 0.834 |
| M8 | s(pds) | 0.01 | 3.00 | 0.00 | 0.9756 | 0.0146 | 7 | -2724.65 | -2697.61 | 1369.33 | M8 vs. M9 | 1.46 | 0.834 |
| | ti(age) | 0.00 | 3.00 | 0.00 | 1.0000 | | | | | | | | |
| M9 | ti(pds) | 0.00 | 3.00 | 0.00 | 0.4526 | 0.0159 | 11 | -2718.11 | -2675.61 | 1370.06 | | | |
| | ti(age, pds) | 1.01 | 1.01 | 1.45 | 0.2313 | | | | | | | | |
| **CEN** | **Smooth Terms** | | | | | | **Model Fit** | | | | | | |
| | **Terms** | **edf** | **Ref.df** | **F** | **p-value** | **R2** | **df** | **AIC** | **BIC** | **logLik** | **Test** | **L.Ratio** | **p-value** |
| M7 | s(age) | 0.84 | 3.00 | 0.88 | 0.0647 | 0.0479 | 7 | -3369.94 | -3342.90 | 1691.97 | M7 vs. M9 | 1.66 | 0.797 |
| M8 | s(pds) | 0.35 | 3.00 | 0.18 | 0.2174 | 0.0422 | 7 | -3368.77 | -3341.72 | 1691.38 | M8 vs. M9 | 2.84 | 0.585 |
| | ti(age) | 0.98 | 3.00 | 1.62 | 0.0174 | | | | | | | | |
| M9 | ti(pds) | 0.01 | 3.00 | 0.00 | 0.6373 | 0.0587 | 11 | -3363.61 | -3321.11 | 1692.80 | | | |
| | ti(age, pds) | 1.95 | 1.95 | 2.18 | 0.1523 | | | | | | | | |
| **AAA** | **Smooth Terms** | | | | | | **Model Fit** | | | | | | |
| | **Terms** | **edf** | **Ref.df** | **F** | **p-value** | **R2** | **df** | **AIC** | **BIC** | **logLik** | **Test** | **L.Ratio** | **p-value** |
| M7 | s(age) | 0.16 | 3.00 | 0.06 | 0.2870 | 0.0064 | 7 | -3206.25 | -3179.20 | 1610.12 | M7 vs. M9 | 3.28 | 0.513 |
| M8 | s(pds) | 0.69 | 3.00 | 0.69 | 0.0840 | 0.0117 | 7 | -3207.15 | -3180.10 | 1610.57 | M8 vs. M9 | 2.37 | 0.667 |
| | ti(age) | 0.00 | 3.00 | 0.00 | 0.6156 | | | | | | | | |
| M9 | ti(pds) | 0.00 | 3.00 | 0.00 | 0.3275 | 0.0123 | 11 | -3201.52 | -3159.02 | 1611.76 | | | |
| | ti(age, pds) | 1.00 | 1.00 | 3.24 | 0.0721 | | | | | | | | |
| **ATA** | **Smooth Terms** | | | | | | **Model Fit** | | | | | | |
| | **Terms** | **edf** | **Ref.df** | **F** | **p-value** | **R2** | **df** | **AIC** | **BIC** | **logLik** | **Test** | **L.Ratio** | **p-value** |
| M7 | s(age) | 2.23 | 3.00 | 2.59 | 0.0222 | 0.0728 | 7 | -2874.39 | -2847.34 | 1444.19 | M7 vs. M9 | 6.70 | 0.153 |
| M8 | s(pds) | 0.70 | 3.00 | 0.63 | 0.0978 | 0.0575 | 7 | -2874.21 | -2847.17 | 1444.11 | M8 vs. M9 | 6.87 | 0.143 |
| | ti(age) | 0.00 | 3.00 | 0.00 | 0.5422 | | | | | | | | |
| M9 | ti(pds) | 0.00 | 3.00 | 0.00 | 0.7251 | 0.0744 | 11 | -2873.09 | -2830.59 | 1447.54 | | | |
| | ti(age, pds) | 1.76 | 1.76 | 2.95 | 0.0324 | | | | | | | | |
| **ASTA** | **Smooth Terms** | | | | | | **Model Fit** | | | | | | |
| | **Terms** | **edf** | **Ref.df** | **F** | **p-value** | **R2** | **df** | **AIC** | **BIC** | **logLik** | **Test** | **L.Ratio** | **p-value** |
| M7 | s(age) | 0.02 | 3.00 | 0.01 | 0.3529 | 0.0161 | 7 | -3073.28 | -3046.24 | 1543.64 | M7 vs. M9 | 0.05 | 1.000 |
| M8 | s(pds) | 0.00 | 3.00 | 0.00 | 0.5596 | 0.0160 | 7 | -3073.28 | -3046.24 | 1543.64 | M8 vs. M9 | 0.05 | 1.000 |
| | ti(age) | 0.00 | 3.00 | 0.00 | 0.3526 | | | | | | | | |
| M9 | ti(pds) | 0.00 | 3.00 | 0.00 | 0.5777 | 0.0133 | 11 | -3065.33 | -3022.83 | 1543.66 | | | |
| | ti(age, pds) | 1.00 | 1.00 | 0.05 | 0.8276 | | | | | | | | |

Notes: In each model, the smooth terms, the estimated degree of freedom (edf), reference degree of freedom (Ref.df), F-score, p-value, and Adjusted $R^2$ for each model is shown; p-value < 0.0056 bolded (Bonferroni corrected). Between model comparisons include the df, AIC, log-likelihood ratio (L Ratio) and p-values < 0.05 bolded.

**Supporting Table 5.** *GAMM amygdala subregion Relative Volume Fraction (RVF) controlling for Intracranial Volume (ICV) results for age, pubertal status, and age-by-pubertal status interaction for males*

| ***MALES*** | | | | | | | | | | | | | |
|---|---|---|---|---|---|---|---|---|---|---|---|---|---|
| **LA** | **Smooth Terms** | | | | | | **Model Fit** | | | | | | |
| | **Terms** | **edf** | **Ref.df** | **F** | **p-value** | **R2** | **df** | **AIC** | **BIC** | **logLik** | **Test** | **L.Ratio** | **p-value** |
| SM2 | s(age) | 1.23 | 3.00 | 2.34 | 0.0068 | 0.0743 | 8 | -2778.28 | -2745.27 | 1397.14 | SM2 vs. SM4 | 0.30 | 0.9899 |
| SM3 | s(pds) | 0.90 | 3.00 | 1.31 | 0.0299 | 0.0647 | 8 | -2776.08 | -2743.07 | 1396.04 | SM3 vs. SM4 | 2.50 | 0.6452 |
| SM4 | ti(age) | 0.95 | 3.00 | 1.78 | 0.0132 | 0.0721 | 12 | -2770.58 | -2721.06 | 1397.29 | | | |
| | ti(pds) | 0.00 | 3.00 | 0.00 | 0.6095 | | | | | | | | |
| | ti(age, pds) | 1.00 | 1.00 | 0.35 | 0.5555 | | | | | | | | |
| **BLDI** | **Smooth Terms** | | | | | | **Model Fit** | | | | | | |
| | **Terms** | **edf** | **Ref.df** | **F** | **p-value** | **R2** | **df** | **AIC** | **BIC** | **logLik** | **Test** | **L.Ratio** | **p-value** |
| SM2 | s(age) | 0.49 | 3.00 | 0.25 | 0.2186 | 0.1373 | 8 | -3875.08 | -3842.06 | 1945.54 | SM2 vs. SM4 | 1.65 | 0.7997 |
| SM3 | s(pds) | 0.80 | 3.00 | 0.95 | 0.0532 | 0.1436 | 8 | -3876.45 | -3843.44 | 1946.23 | SM3 vs. SM4 | 0.28 | 0.9913 |
| SM4 | ti(age) | 0.00 | 3.00 | 0.00 | 0.8283 | 0.1422 | 12 | -3868.73 | -3819.21 | 1946.36 | | | |
| | ti(pds) | 0.69 | 3.00 | 0.71 | 0.0762 | | | | | | | | |
| | ti(age, pds) | 1.01 | 1.01 | 0.29 | 0.5932 | | | | | | | | |
| **BLVPL** | **Smooth Terms** | | | | | | **Model Fit** | | | | | | |
| | **Terms** | **edf** | **Ref.df** | **F** | **p-value** | **R2** | **df** | **AIC** | **BIC** | **logLik** | **Test** | **L.Ratio** | **p-value** |
| SM2 | s(age) | 2.37 | 3.00 | 10.98 | ***5.97E-08*** | 0.1415 | 8 | -3628.90 | -3595.88 | 1822.45 | SM2 vs. SM4 | 2.78 | 0.5946 |
| SM3 | s(pds) | 1.12 | 3.00 | 2.99 | ***0.0021*** | 0.1069 | 8 | -3676.47 | -3643.45 | 1846.23 | SM3 vs. SM4 | 44.78 | ***4.41E-09*** |
| SM4 | ti(age) | 2.05 | 3.00 | 6.68 | ***3.85E-06*** | 0.1737 | 12 | -3623.68 | -3574.16 | 1823.84 | | | |
| | ti(pds) | 0.01 | 3.00 | 0.00 | 0.2716 | | | | | | | | |
| | ti(age, pds) | 5.65 | 5.65 | 2.26 | 0.1523 | | | | | | | | |
| **BM** | **Smooth Terms** | | | | | | **Model Fit** | | | | | | |
| | **Terms** | **edf** | **Ref.df** | **F** | **p-value** | **R2** | **df** | **AIC** | **BIC** | **logLik** | **Test** | **L.Ratio** | **p-value** |
| SM2 | s(age) | 0.04 | 3.00 | 0.01 | 0.4043 | 0.0592 | 8 | -3872.27 | -3839.25 | 1944.13 | SM2 vs. SM4 | 0.67 | 0.9544 |
| SM3 | s(pds) | 0.32 | 3.00 | 0.15 | 0.2422 | 0.0600 | 8 | -3872.34 | -3839.33 | 1944.17 | SM3 vs. SM4 | 0.60 | 0.9628 |
| SM4 | ti(age) | 0.01 | 3.00 | 0.00 | 0.5908 | 0.0639 | 12 | -3864.94 | -3815.42 | 1944.47 | | | |
| | ti(pds) | 0.03 | 3.00 | 0.01 | 0.3782 | | | | | | | | |
| | ti(age, pds) | 2.46 | 2.46 | 0.44 | 0.5577 | | | | | | | | |
| **CMN** | **Smooth Terms** | | | | | | **Model Fit** | | | | | | |
| | **Terms** | **edf** | **Ref.df** | **F** | **p-value** | **R2** | **df** | **AIC** | **BIC** | **logLik** | **Test** | **L.Ratio** | **p-value** |
| SM2 | s(age) | 0.10 | 3.00 | 0.03 | 0.3288 | 0.0199 | 8 | -3500.73 | -3467.71 | 1758.36 | SM2 vs. SM4 | 0.30 | 0.9899 |
| SM3 | s(pds) | 0.02 | 3.00 | 0.00 | 0.7721 | 0.0195 | 8 | -3500.72 | -3467.70 | 1758.36 | SM3 vs. SM4 | 0.31 | 0.9892 |
| SM4 | ti(age) | 0.00 | 3.00 | 0.00 | 0.3778 | 0.0186 | 12 | -3493.03 | -3443.50 | 1758.51 | | | |
| | ti(pds) | 0.00 | 3.00 | 0.00 | 0.9150 | | | | | | | | |
| | ti(age, pds) | 1.06 | 1.06 | 0.26 | 0.6062 | | | | | | | | |
| **CEN** | **Smooth Terms** | | | | | | **Model Fit** | | | | | | |
| | **Terms** | **edf** | **Ref.df** | **F** | **p-value** | **R2** | **df** | **AIC** | **BIC** | **logLik** | **Test** | **L.Ratio** | **p-value** |
| SM2 | s(age) | 2.51 | 3.00 | 10.92 | ***1.04E-07*** | 0.0884 | 8 | -4273.01 | -4239.99 | 2144.50 | SM2 vs. SM4 | 0.62 | 0.9608 |
| SM3 | s(pds) | 1.10 | 3.00 | 2.95 | ***0.0021*** | 0.0516 | 8 | -4322.06 | -4289.05 | 2169.03 | SM3 vs. SM4 | 48.43 | ***7.66E-10*** |
| SM4 | ti(age) | 2.48 | 3.00 | 9.72 | ***1.82E-07*** | 0.0909 | 12 | -4265.63 | -4216.11 | 2144.81 | | | |
| | ti(pds) | 0.00 | 3.00 | 0.00 | 0.8121 | | | | | | | | |
| | ti(age, pds) | 1.29 | 1.29 | 0.63 | 0.5183 | | | | | | | | |
| **AAA** | **Smooth Terms** | | | | | | **Model Fit** | | | | | | |
| | **Terms** | **edf** | **Ref.df** | **F** | **p-value** | **R2** | **df** | **AIC** | **BIC** | **logLik** | **Test** | **L.Ratio** | **p-value** |
| SM2 | s(age) | 0.01 | 3.00 | 0.00 | 0.5100 | -0.0063 | 8 | -4119.80 | -4086.78 | 2067.90 | SM2 vs. SM4 | 2.00 | 0.7353 |
| SM3 | s(pds) | 0.01 | 3.00 | 0.00 | 0.9769 | -0.0063 | 8 | -4119.79 | -4086.78 | 2067.90 | SM3 vs. SM4 | 2.00 | 0.7350 |
| SM4 | ti(age) | 0.01 | 3.00 | 0.00 | 0.5126 | 0.0097 | 12 | -4113.80 | -4064.28 | 2068.90 | | | |
| | ti(pds) | 0.01 | 3.00 | 0.00 | 0.9057 | | | | | | | | |
| | ti(age, pds) | 3.75 | 3.75 | 1.91 | 0.1546 | | | | | | | | |
| **ATA** | **Smooth Terms** | | | | | | **Model Fit** | | | | | | |
| | **Terms** | **edf** | **Ref.df** | **F** | **p-value** | **R2** | **df** | **AIC** | **BIC** | **logLik** | **Test** | **L.Ratio** | **p-value** |
| SM2 | s(age) | 2.47 | 3.00 | 22.93 | ***1.83E-15*** | 0.1546 | 8 | -3671.26 | -3638.24 | 1843.63 | SM2 vs. SM4 | 59.43 | ***3.83E-12*** |
| SM3 | s(pds) | 1.32 | 3.00 | 5.61 | ***3.41E-05*** | 0.0805 | 8 | -3709.19 | -3676.18 | 1862.60 | SM3 vs. SM4 | 21.49 | ***0.0003*** |
| SM4 | ti(age) | 2.16 | 3.00 | 10.78 | ***2.37E-08*** | 0.1537 | 12 | -3722.68 | -3673.16 | 1873.34 | | | |
| | ti(pds) | 0.00 | 3.00 | 0.00 | 0.5004 | | | | | | | | |
| | ti(age, pds) | 1.01 | 1.01 | 0.04 | 0.8486 | | | | | | | | |
| **ASTA** | **Smooth Terms** | | | | | | **Model Fit** | | | | | | |
| | **Terms** | **edf** | **Ref.df** | **F** | **p-value** | **R2** | **df** | **AIC** | **BIC** | **logLik** | **Test** | **L.Ratio** | **p-value** |
| SM2 | s(age) | 1.63 | 3.00 | 4.17 | ***0.0006*** | 0.0347 | 8 | -3980.75 | -3947.73 | 1998.37 | SM2 vs. SM4 | 2.12 | 0.7144 |
| SM3 | s(pds) | 0.91 | 3.00 | 1.38 | 0.0264 | 0.0172 | 8 | -3974.75 | -3941.73 | 1995.37 | SM3 vs. SM4 | 8.11 | 0.0875 |
| SM4 | ti(age) | 0.79 | 3.00 | 1.04 | 0.0343 | 0.0409 | 12 | -3974.86 | -3925.34 | 1999.43 | | | |
| | ti(pds) | 0.01 | 3.00 | 0.00 | 0.4430 | | | | | | | | |
| | ti(age, pds) | 2.87 | 2.87 | 2.24 | 0.1516 | | | | | | | | |

Notes: In each model, the smooth terms, the estimated degree of freedom (edf), reference degree of freedom (Ref.df), F-score, p-value, and Adjusted $R^2$ for each model is shown; p-value < 0.0056 bolded (Bonferroni corrected). Between model comparisons include the df, AIC, log-likelihood ratio (L Ratio) and p-values < 0.05 bolded.

**Supplemental Table 6.** *GAMM amygdala subregion Relative Volume Fraction (RVF) controlling for Intracranial Volume (ICV) results for age, pubertal status, and age-by-pubertal status interaction for females*

| | | | | | | ***FEMALES*** | | | | | | | |
|---|---|---|---|---|---|---|---|---|---|---|---|---|---|
| **LA** | **Smooth Terms** | | | | | | **Model Fit** | | | | | | |
| | **Terms** | **edf** | **Ref.df** | **F** | **p-value** | **R2** | **df** | **AIC** | **BIC** | **logLik** | **Test** | **L.Ratio** | **p-value** |
| SM2 | s(age) | 0.22 | 3.00 | 0.09 | 0.2706 | 0.0361 | 8 | -2167.68 | -2136.77 | 1091.84 | SM2 vs. SM4 | 2.20 | 0.699 |
| SM3 | s(pds) | 0.27 | 3.00 | 0.13 | 0.2442 | 0.0364 | 8 | -2167.70 | -2136.80 | 1091.85 | SM3 vs. SM4 | 2.18 | 0.703 |
| SM4 | ti(age) | 0.00 | 3.00 | 0.00 | 0.5007 | 0.0387 | 12 | -2161.88 | -2115.52 | 1092.94 | | | |
| | ti(pds) | 0.00 | 3.00 | 0.00 | 0.6377 | | | | | | | | |
| | ti(age, pds) | 1.00 | 1.00 | 2.20 | 0.1387 | | | | | | | | |
| **BLDI** | **Smooth Terms** | | | | | | **Model Fit** | | | | | | |
| | **Terms** | **edf** | **Ref.df** | **F** | **p-value** | **R2** | **df** | **AIC** | **BIC** | **logLik** | **Test** | **L.Ratio** | **p-value** |
| SM2 | s(age) | 0.02 | 3.00 | 0.00 | 1.0000 | 0.1807 | 8 | -3019.22 | -2988.31 | 1517.61 | SM2 vs. SM4 | 0.16 | 0.997 |
| SM3 | s(pds) | 0.02 | 3.00 | 0.00 | 1.0000 | 0.1807 | 8 | -3019.22 | -2988.31 | 1517.61 | SM3 vs. SM4 | 0.16 | 0.997 |
| SM4 | ti(age) | 0.00 | 3.00 | 0.00 | 1.0000 | 0.1789 | 12 | -3011.38 | -2965.02 | 1517.69 | | | |
| | ti(pds) | 0.00 | 3.00 | 0.00 | 0.9335 | | | | | | | | |
| | ti(age, pds) | 1.00 | 1.00 | 0.15 | 0.7020 | | | | | | | | |
| **BLVPL** | **Smooth Terms** | | | | | | **Model Fit** | | | | | | |
| | **Terms** | **edf** | **Ref.df** | **F** | **p-value** | **R2** | **df** | **AIC** | **BIC** | **logLik** | **Test** | **L.Ratio** | **p-value** |
| SM2 | s(age) | 0.00 | 3.00 | 0.00 | 0.6675 | 0.0871 | 8 | -2872.72 | -2841.81 | 1444.36 | SM2 vs. SM4 | 3.37 | 0.498 |
| SM3 | s(pds) | 0.02 | 3.00 | 0.01 | 0.3207 | 0.0871 | 8 | -2872.73 | -2841.82 | 1444.36 | SM3 vs. SM4 | 3.37 | 0.498 |
| SM4 | ti(age) | 1.85 | 3.00 | 2.24 | 0.0151 | 0.1279 | 12 | -2868.09 | -2821.73 | 1446.05 | | | |
| | ti(pds) | 0.00 | 3.00 | 0.00 | 0.8469 | | | | | | | | |
| | ti(age, pds) | 3.84 | 3.84 | 2.72 | 0.0468 | | | | | | | | |
| **BM** | **Smooth Terms** | | | | | | **Model Fit** | | | | | | |
| | **Terms** | **edf** | **Ref.df** | **F** | **p-value** | **R2** | **df** | **AIC** | **BIC** | **logLik** | **Test** | **L.Ratio** | **p-value** |
| SM2 | s(age) | 0.01 | 3.00 | 0.00 | 0.4809 | 0.0474 | 8 | -3126.19 | -3095.28 | 1571.10 | SM2 vs. SM4 | 1.63 | 0.804 |
| SM3 | s(pds) | 0.04 | 3.00 | 0.02 | 0.3141 | 0.0475 | 8 | -3126.20 | -3095.29 | 1571.10 | SM3 vs. SM4 | 1.62 | 0.804 |
| SM4 | ti(age) | 0.00 | 3.00 | 0.00 | 0.7273 | 0.0490 | 12 | -3119.82 | -3073.46 | 1571.91 | | | |
| | ti(pds) | 0.00 | 3.00 | 0.00 | 0.6708 | | | | | | | | |
| | ti(age, pds) | 1.01 | 1.01 | 1.61 | 0.2065 | | | | | | | | |
| **CMN** | **Smooth Terms** | | | | | | **Model Fit** | | | | | | |
| | **Terms** | **edf** | **Ref.df** | **F** | **p-value** | **R2** | **df** | **AIC** | **BIC** | **logLik** | **Test** | **L.Ratio** | **p-value** |
| SM2 | s(age) | 0.00 | 3.00 | 0.00 | 0.6658 | 0.0367 | 8 | -2731.67 | -2700.76 | 1373.84 | SM2 vs. SM4 | 1.46 | 0.834 |
| SM3 | s(pds) | 0.00 | 3.00 | 0.00 | 0.9366 | 0.0367 | 8 | -2731.67 | -2700.76 | 1373.84 | SM3 vs. SM4 | 1.46 | 0.834 |
| SM4 | ti(age) | 0.00 | 3.00 | 0.00 | 0.9994 | 0.0380 | 12 | -2725.13 | -2678.77 | 1374.57 | | | |
| | ti(pds) | 0.00 | 3.00 | 0.00 | 0.5069 | | | | | | | | |
| | ti(age, pds) | 1.01 | 1.01 | 1.45 | 0.2311 | | | | | | | | |
| **CEN** | **Smooth Terms** | | | | | | **Model Fit** | | | | | | |
| | **Terms** | **edf** | **Ref.df** | **F** | **p-value** | **R2** | **df** | **AIC** | **BIC** | **logLik** | **Test** | **L.Ratio** | **p-value** |
| SM2 | s(age) | 0.83 | 3.00 | 0.87 | 0.0661 | 0.0452 | 8 | -3367.97 | -3337.06 | 1691.99 | SM2 vs. SM4 | 1.66 | 0.799 |
| SM3 | s(pds) | 0.34 | 3.00 | 0.18 | 0.2205 | 0.0395 | 8 | -3366.80 | -3335.90 | 1691.40 | SM3 vs. SM4 | 2.82 | 0.588 |
| SM4 | ti(age) | 0.98 | 3.00 | 1.60 | 0.0181 | 0.0560 | 12 | -3361.63 | -3315.26 | 1692.81 | | | |
| | ti(pds) | 0.01 | 3.00 | 0.00 | 0.6388 | | | | | | | | |
| | ti(age, pds) | 1.95 | 1.95 | 2.17 | 0.1549 | | | | | | | | |
| **AAA** | **Smooth Terms** | | | | | | **Model Fit** | | | | | | |
| | **Terms** | **edf** | **Ref.df** | **F** | **p-value** | **R2** | **df** | **AIC** | **BIC** | **logLik** | **Test** | **L.Ratio** | **p-value** |
| SM2 | s(age) | 0.13 | 3.00 | 0.05 | 0.2964 | 0.0142 | 8 | -3208.06 | -3177.15 | 1612.03 | SM2 vs. SM4 | 3.35 | 0.501 |
| SM3 | s(pds) | 0.67 | 3.00 | 0.64 | 0.0925 | 0.0191 | 8 | -3208.88 | -3177.97 | 1612.44 | SM3 vs. SM4 | 2.54 | 0.638 |
| SM4 | ti(age) | 0.00 | 3.00 | 0.00 | 0.6406 | 0.0203 | 12 | -3203.41 | -3157.05 | 1613.71 | | | |
| | ti(pds) | 0.00 | 3.00 | 0.00 | 0.3564 | | | | | | | | |
| | ti(age, pds) | 1.00 | 1.00 | 3.30 | 0.0696 | | | | | | | | |
| **ATA** | **Smooth Terms** | | | | | | **Model Fit** | | | | | | |
| | **Terms** | **edf** | **Ref.df** | **F** | **p-value** | **R2** | **df** | **AIC** | **BIC** | **logLik** | **Test** | **L.Ratio** | **p-value** |
| SM2 | s(age) | 2.11 | 3.00 | 2.22 | 0.0351 | 0.0790 | 8 | -2876.40 | -2845.49 | 1446.20 | SM2 vs. SM4 | 7.19 | 0.126 |
| SM3 | s(pds) | 0.73 | 3.00 | 0.73 | 0.0847 | 0.0675 | 8 | -2876.85 | -2845.94 | 1446.42 | SM3 vs. SM4 | 6.74 | 0.1502 |
| SM4 | ti(age) | 0.00 | 3.00 | 0.00 | 0.5821 | 0.0834 | 12 | -2875.59 | -2829.23 | 1449.79 | | | |
| | ti(pds) | 0.00 | 3.00 | 0.00 | 0.8022 | | | | | | | | |
| | ti(age, pds) | 1.74 | 1.74 | 2.91 | 0.0338 | | | | | | | | |
| **ASTA** | **Smooth Terms** | | | | | | **Model Fit** | | | | | | |
| | **Terms** | **edf** | **Ref.df** | **F** | **p-value** | **R2** | **df** | **AIC** | **BIC** | **logLik** | **Test** | **L.Ratio** | **p-value** |
| SM2 | s(age) | 0.02 | 3.00 | 0.00 | 0.3632 | 0.0165 | 8 | -3072.47 | -3041.56 | 1544.23 | SM2 vs. SM4 | 0.05 | 1.000 |
| SM3 | s(pds) | 0.00 | 3.00 | 0.00 | 0.5791 | 0.0165 | 8 | -3072.47 | -3041.56 | 1544.23 | SM3 vs. SM4 | 0.05 | 1.000 |
| SM4 | ti(age) | 0.00 | 3.00 | 0.00 | 0.3642 | 0.0138 | 12 | -3064.52 | -3018.15 | 1544.26 | | | |
| | ti(pds) | 0.00 | 3.00 | 0.00 | 0.6014 | | | | | | | | |
| | ti(age, pds) | 1.00 | 1.00 | 0.05 | 0.8235 | | | | | | | | |

Notes: In each model, the smooth terms, the estimated degree of freedom (edf), reference degree of freedom (Ref.df), F-score, p-value, and Adjusted $R^2$ for each model is shown; p-value < 0.0056 bolded (Bonferroni corrected). Between model comparisons include the df, AIC, log-likelihood ratio (L Ratio) and p-values < 0.05 bolded.

**Supporting Information**

**Figures**

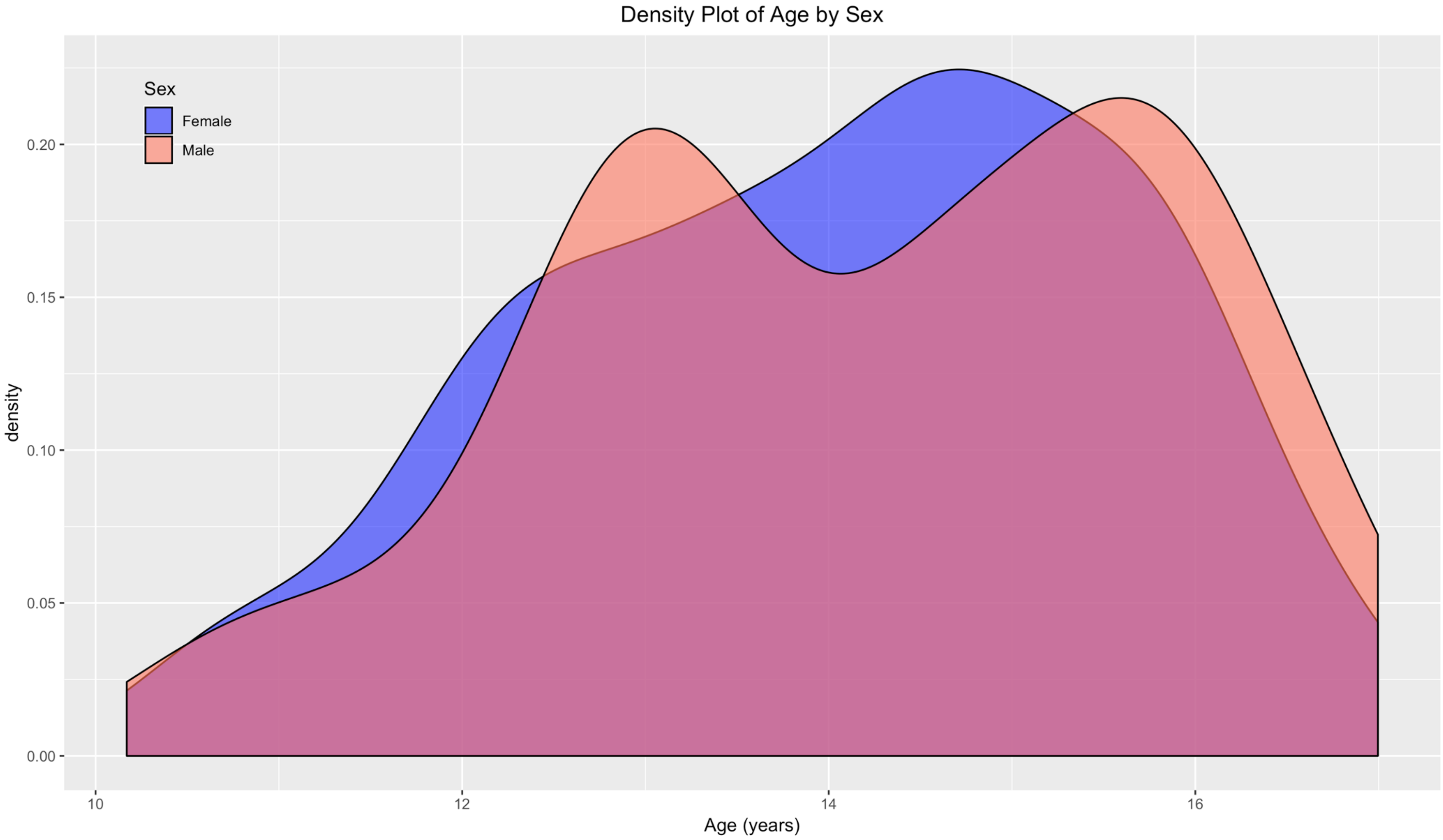


**Supporting Figure 1: Density plot of age distribution by sex**. Age is a numeric variable based on the difference in years from the date of visit and date of birth of subject to two decimal points.

**Supporting Information**

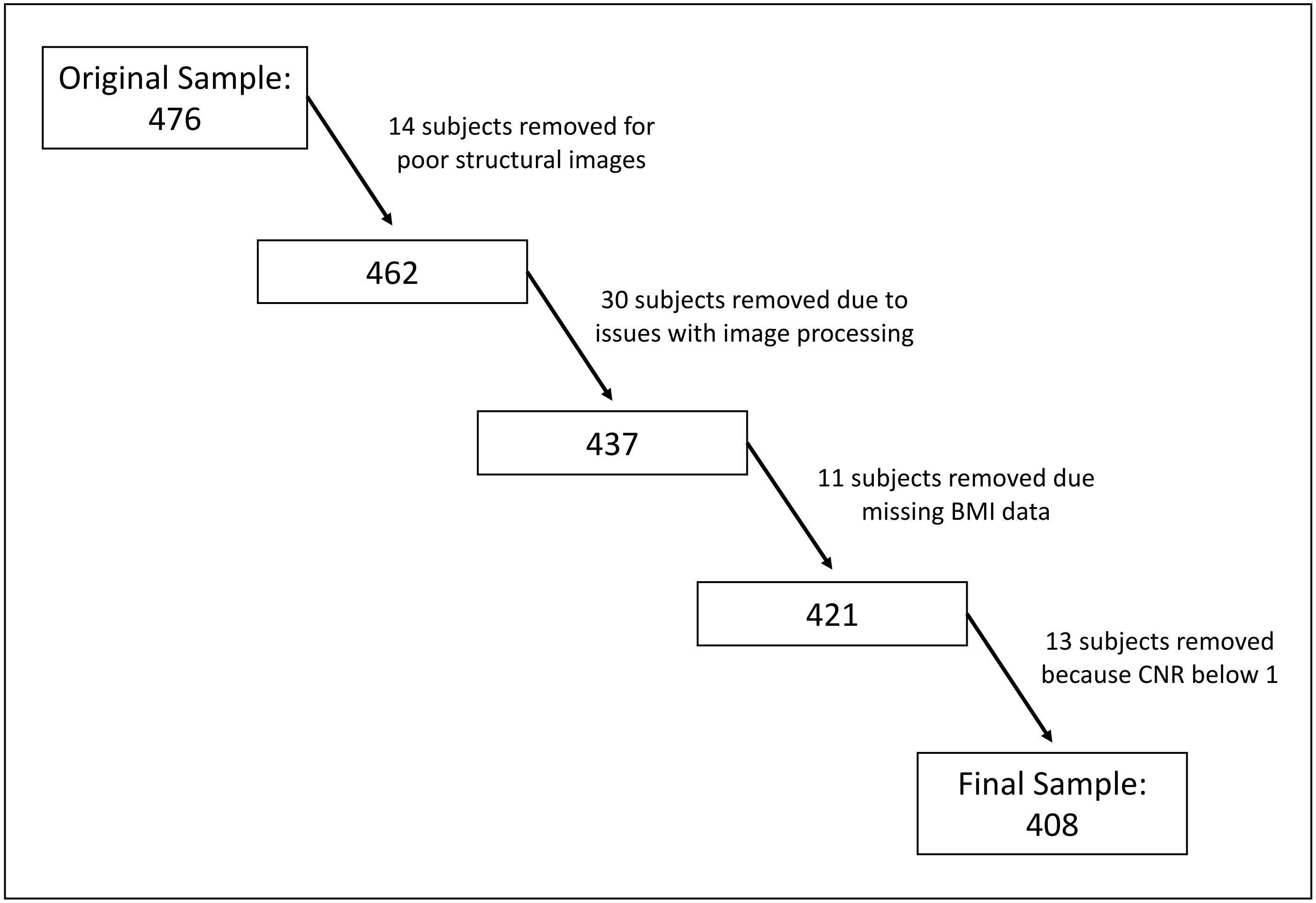


**Supporting Figure 2: Flowchart of QC and selection of subjects**. Final sample of 408 demographics can be seen in Table 1.

**Supporting Information**

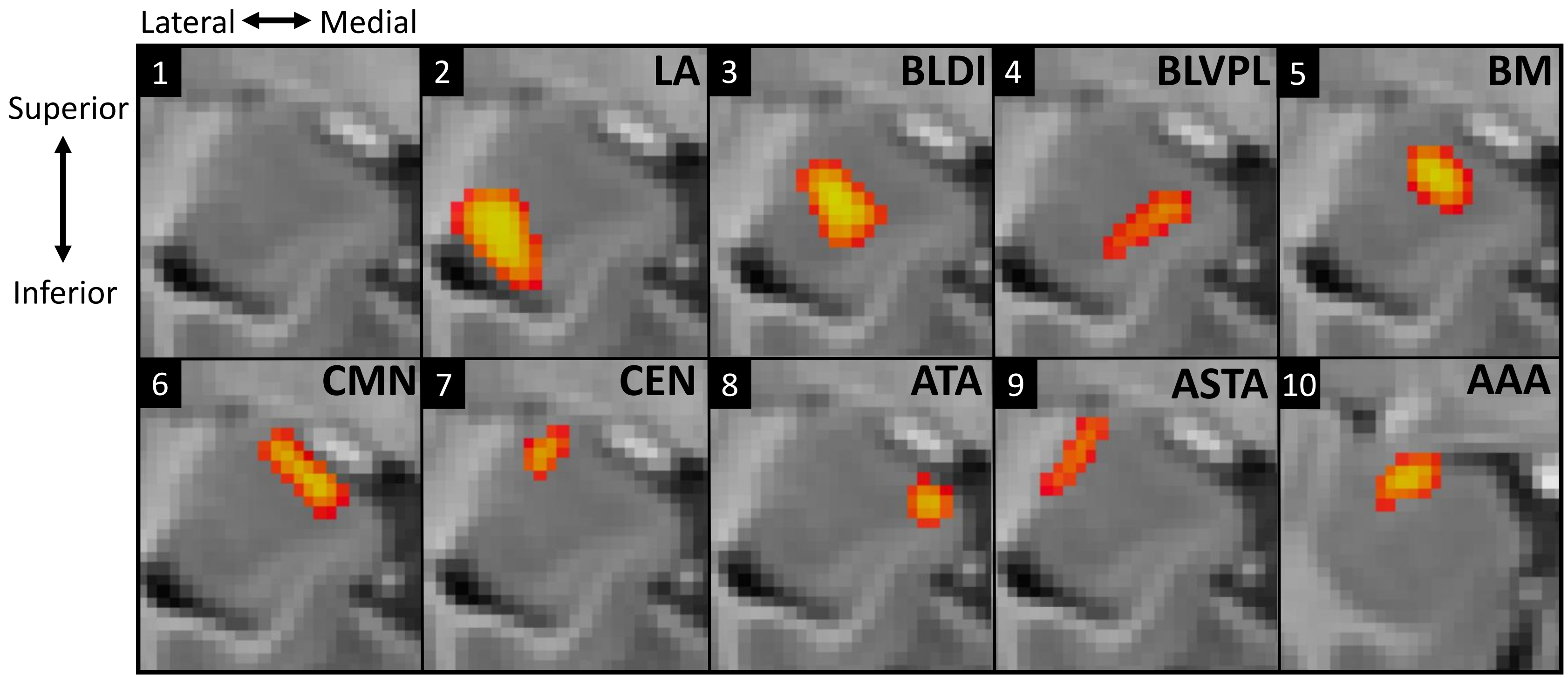


**Supporting Figure 3: Group level amygdala segmentation on a minimum deformation template (MDT) of 10 to 12-year-old females in our sample.** Each block is oriented in the same orientation as block 1; block 1 has no segmentation, the following segmentations are labeled as follows: lateral (LA), dorsal and intermediate divisions of the basolateral nucleus (BLDI), ventral division of the basolateral nucleus and paralaminar nucleus (BLVPL), basomedial nucleus (BM), cortical and medial nuclei (CMN), central nucleus (CEN), amygdala transition areas (ATA), amygdalostriatal transition area (ASTA), anterior amygdala area (AAA).

**Supporting Information**

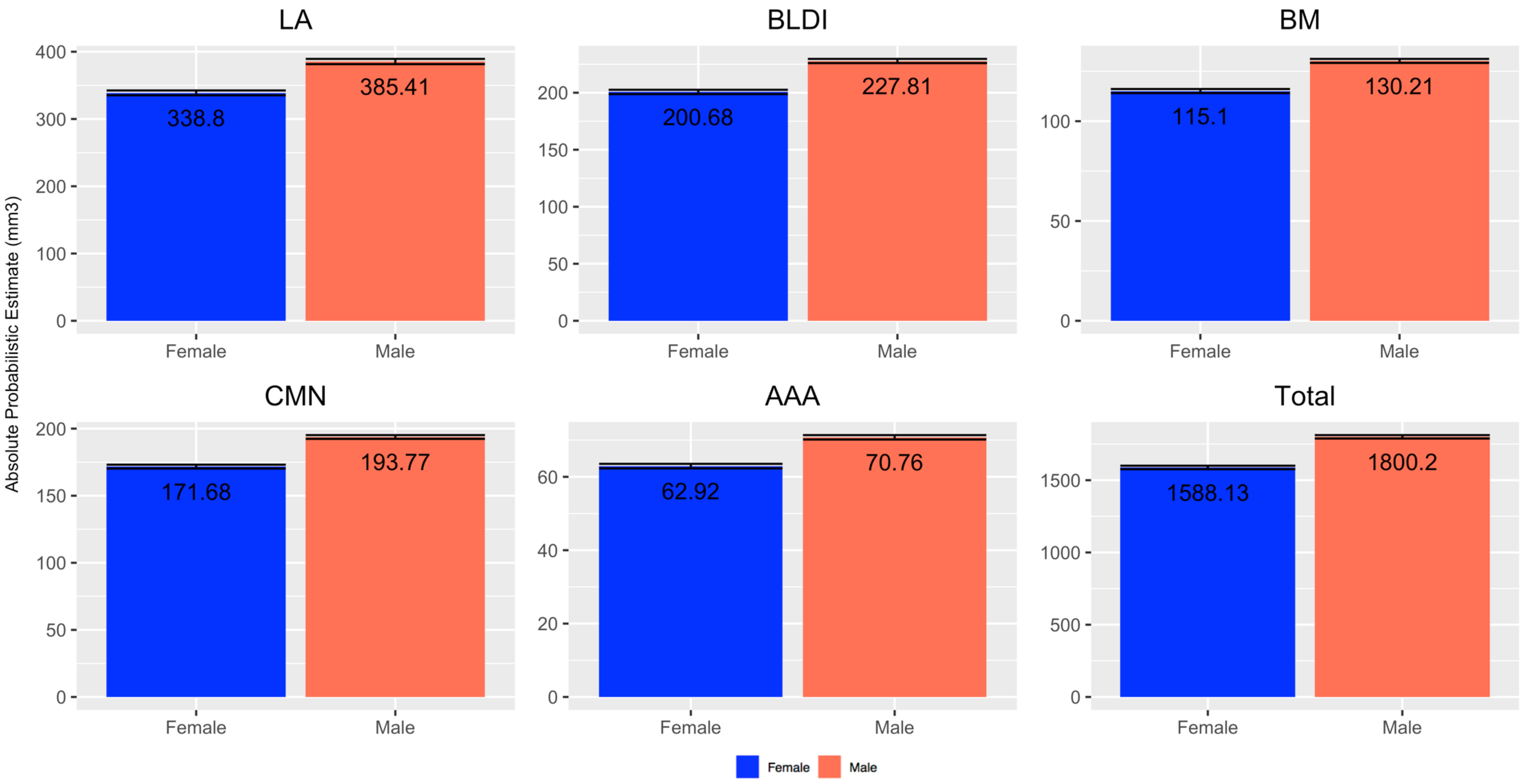


**Supporting Figure 4: Bar graph of main sex effect of amygdala subregions**. Means and standard error bars of the absolute probabilistic estimate ($mm^3$) by sex graphed for lateral (LA), basolateral dorsal and intermediate subdivision (BLDI), basomedial nucleus (BM), cortical and medial nuclei (CMN), anterior amygdala area (AAA), and total amygdala volume (Total).
